# A polyomavirus-positive Merkel cell carcinoma mouse model supports a unified cancer origin

**DOI:** 10.1101/2025.05.06.652325

**Authors:** Wendy Yang, Sara Contente, Sarah Rahman

## Abstract

The Germ Cell Theory, rooted in developmental biology, has significantly advanced both the understanding and curative treatment of rare germ cell cancers (GCCs). In contrast, somatic cancer research - long dominated by the Somatic Mutation Theory - has stagnated due to fragmented, mutation-centric approaches that lack a unifying conceptual framework and have yielded only limited clinical progress. GCC research identifies non-somatic human primordial germ cells (hPGCs) as the cells of origin for GCCs. In somatic cancers, although the cancer stem cell theory is gaining acceptance, cancer stem cells are assumed to be of somatic origin, with their exact identity undefined. Accumulating experimental evidence and clinical observations challenge the traditional separation between GCCs and somatic cancers, including malignant somatic transformation (MST) - the emergence of somatic cancer phenotypes from GCCs, often without new mutations - suggesting that hPGCs may also initiate somatic cancers. However, no experimental model of MST has been established. Merkel cell polyomavirus-positive Merkel cell carcinoma (MCC), a highly aggressive somatic cancer with paradoxically stable genomes resembling those of GCCs, offers a unique model to test germ cell theory-based somatic oncogenesis. Here we present an MST mouse model linking hPGC origin to somatic cancer by inducing virus-positive MCC-like tumors in vivo from virus-transfected hPGC-like cells or human iPSCs competent for hPGC specification with an obligatory late-hPGC state preceding MST. This genetically simple, molecularly tractable model provides a novel platform to dissect VP-MCC pathogenesis and broadly advocates a developmental biology framework for somatic cancer research beyond mutation-centric and soma-centric paradigms.

## Introduction

Cancer has traditionally been classified into two primary groups based on presumed cellular origin: germ cell cancers (GCCs), arising from reproductive germline cells - also referred to as Type II germ cell tumors (GCTs) - which constitute less than 1% of malignancies (Oosterhuis et al., 2019); and somatic cancers, originating from non-germline somatic cells, representing the vast majority of malignancies.

Despite this traditional separation, all cancers share phenotypic hallmarks, many of which recapitulate early embryonic/germ cell development, suggesting developmental arrest or “blocked ontogeny” (Hanahan et al., 2011; Sell et al., 1994). This principle, encapsulated as “oncology recapitulates ontogeny,” has significantly influenced GCC research and clinical practice yet remains largely overlooked in somatic cancer studies (Tu et al., 2021; Huang et al., 2025), despite its clinical relevance to grading and prognosis (Jögi et al., 2012).

The Germ Cell Theory, grounded in developmental biology, has yielded transformative insights into GCT biology and curative treatment strategies (Oosterhuis et al., 2019; Tu et al., 2021). GCCs are genetically simple and epigenetically distinct (Oosterhuis et al., 2019; Tu et al., 2021), originating from human primordial germ cells (hPGCs) (Oosterhuis et al., 2019; Taguch et al., 2021) - founder cells of the germline specified during weeks 2-3 of development (Chen, D. *et al*. 2019). hPGCs exhibit latent pluripotency and intrinsic cancer stem cell-like properties, predisposing them to malignant transformation with minimal genetic alteration (Oosterhuis et al., 2019). In GCCs, transformed hPGCs and their totipotent derivatives - embryonal carcinoma (EC) cells thought to arise via parthenogenesis in vivo - constitute the cancer stem cell compartments of seminomas and non-seminomatous germ cell cancers (NSGCCs), respectively (Oosterhuis et al., 2019). Seminomas are composed entirely of transformed hPGCs, while NSGCCs such as ECs or malignant teratomas consist solely of EC cells or arise through multilineage somatic differentiation from ECs respectively (Oosterhuis et al., 2019).

Mounting biological and clinical evidence increasingly challenges the traditional separation between germ cell cancers (GCCs) and somatic cancers, suggesting a shared developmental origin. Notably, approximately 3–6% of germ cell tumors (GCTs) undergo malignant somatic transformation (MST) into somatic cancer phenotypes, often without additional mutations (Tu et al., 2021; Kum et al., 2012; Taylor et al., 2020; Wyvekens et al., 2022). GCTs can also manifest as extragonadal GCTs (EGGCTs), demonstrating - like somatic cancers - the capacity to arise at virtually any anatomical site, albeit with a predilection for midline locations (De Felici et al., 2021). Many somatic cancers aberrantly express cancer-testis antigens normally restricted to germ cells (Nin et al., 2023; Simpson et al., 2005), and acquisition of a primordial germ cell (PGC)-like state has been shown to be essential for metastasis (Ma et al., 2023). Furthermore, global DNA hypomethylation is a ubiquitous epigenetic hallmark of all cancers, observed in both GCCs and somatic cancers alike (Oosterhuis et al., 2019; Ehrlich, 2009). Despite these converging features, somatic cancer research remains dominated by the fragmented, mutation-centric Somatic Mutation Theory (SMT), which lacks a unifying biological framework and has yielded limited improvements in patient outcomes (Huang et al., 2025). Although the cancer stem cell theory has more recently gained recognition in somatic cancer research, it fundamentally differs from the germ cell theory in its soma-centric view by positing somatic adult stem cells - whose precise identity remains undefined - as the origin of somatic cancers (Visvader et al., 2008). Given the substantial biological and clinical parallels between GCCs and somatic cancers, it is reasonable to propose that hPGCs - or a PGC-like state - may represent a shared stem cell origin for both. However, no experimental model has yet directly linked hPGCs to somatic cancer via MST (Damjanov et al., 2016), despite the longstanding use of teratoma assays to study GCT development and tumorigenesis (Montilla-Rojo et al., 2023). These assays, which generate teratomas (a form of non-seminomatous GCT) in immunocompromised mice following injection of human pluripotent stem cells (hPSCs), remain the gold standard for confirming pluripotency, due to the intrinsic coupling of pluripotency and teratoma formation (Aleckovic et al., 2008). While MST most commonly arises in clinical teratomas, it has not been recapitulated in these models - likely due to the limited duration of in vivo assays relative to the prolonged latency observed in human disease (Damjanov et al., 2016).

Virus-positive Merkel cell carcinoma (VP-MCC), an aggressive high grade neuroendocrine carcinoma (HGNEC) of the skin, comprises ~80% of MCC cases and is defined by the presence of randomly but clonally integrated, C-terminus truncated Merkel cell polyomavirus (MCPyV) genomes (~5 kb) (Feng et al., 2008; Harms et al., 2018; Pedersen et al., 2024). It presents two key features that seem to make it ideally suited for developing an MST model. First, VP-MCC exhibits extreme genomic stability, in stark contrast to virus-negative MCC (VN-MCC), which comprises the remaining ~20% and is among the most highly mutated human cancers. Similarly, other virus-associated cancers, such as HPV- and EBV-positive tumors, typically show higher mutational burdens than their VN counterparts (Lu et al., 2024). This exceptional genomic stability mirrors that of germ cell tumors (GCTs) and would enable the development of a genetically simple, experimentally tractable MST model. Although MCPyV is ubiquitous in normal skin flora, VP-MCC is very rare, and MCPyV transduction of cutaneous lineages has failed to reproduce its molecular and histologic features - underscoring the paramount importance of cellular context, a principle also similar to GCT biology (Harms et al., 2018; Pedersen et al., 2024; Kervarrec et al., 2019). Second, the small cell neuroendocrine carcinoma (SCNC) phenotype of MCC represents a poorly differentiated, aggressive end-stage state to which diverse somatic cancers converge upon progression (Balanis et al., 2019). SCNCs, including MCC, also exhibit stem-like properties and multilineage differentiation potential (Zhang et al., 2013), suggesting that VP-MCC may be composed predominantly of somatic cancer stem cells, functionally analogous to seminomas or embryonal carcinomas (ECs) in GCTs. Together, these features support the potential of developing a VP-MCC-based MST model capable of capturing transformation events rapidly with immediate somatic differentiation arrests - within the temporal limits of mouse xenograft assays - where MST has otherwise not been observed (Damjanov et al., 2016). To directly test the Germ Cell Theory in somatic cancer, we proceeded to establish a mouse model based on the hypothesis that VP-MCC arises through MCPyV-driven MST of extragonadal hPGCs residing in the adult cutaneous niche. Multiple lines of evidence support our hypothesis. MCPyV tropism toward human primordial germ cells (hPGCs) is indicated by the relatively high level of MCPyV DNA detected in seminoma (Csoboz et al., 2020), a GCC composed of transformed hPGCs (Oosterhuis et al., 2019).

Inactivating mutations of p53 have been associated with MST (Oosterhuis et al., 2019) while MCPyV is known to indirectly inhibit p53 (Pedersen et al., 2024). The skin microenvironment is enriched in stem cell factor (SCF) and stromal cell-derived factor 1 (SDF-1), key regulators of hPGC survival and chemotaxis (Foster et al., 2021; Quan et al., 2015). Furthermore, the male predominance and head and neck predilection of MCC reflect Y chromosome-mediated hPGC viability and their characteristic midline migratory trajectory (Oosterhuis et al., 2019; De Felici et al., 2021).

To investigate the proposed MCPyV-driven VP-MCC tumorigenic pathway, we designed a teratoma assay-like experiment aiming to prove MST from MCPyV transfected hPGCs to VP-MCC like tumors (VMLTs) in the adult mouse cutaneous niche. With the approval of USUHS Institutional Biosafety Committee, we transduced a truncated MCPyV genome (L82) via lentiviral vector into in vitro surrogates of hPGCs and additional two relevant hPSC types - human embryonic germ cells (hEGCs) and human embryonic stem cells (hESCs). Based on prior studies on hPSC-induced mouse teratoma xenografts, we estimated that three mice per condition would provide >99% success rate of forming mouse xenograft tumors (Prokhorova et al., 2009). Eighteen adult NSG mice were divided into three groups (n = 6 per group). The **1**^**st**^ **group** used **hPGC-like cells** (**hPGCLCs**) that are derived from hPSCs and recapitulate early-hPGC (Irie et al., 2015; Sasaki et al., 2015). Both normal hPGCs and hPGCLCs exhibit latent pluripotency - retaining germline unipotency in vivo without chimera (Leitch et al., 2013) or teratoma-forming capacity (Aleckovic et al., 2008; Sugawa et al., 2015) - but can undergo parthenogenetic reversion to pluripotent hEGCs or hEGC-like cells (hEGCLCs) in vitro (Kobayashi et al., 2022; Shamblott et al., 1998). Due to limited culture yield, low-number (~3×10^5^) cells of either hPGCLC_A4 (fig. S1A, provided by the Shioda lab (Kobayashi et al., 2022)) or MCPyV-transduced hPGCLC_A4_L82 (fig. S1B) were injected into the right flanks of three mice per condition. The **2**^**nd**^ **group** used **Human induced pluripotent stem cells** (**hiPSCs**) that are derived from reprogramming of somatic cells and resemble hESCs (Takahashi et al., 2006). Based on our hypothesis, an indirect route from hPSCs to VP-MCC-like tumors (VMLTs) can theoretically be established through hPGCLC specification from competent hPSCs (Irie et al., 2015). Totipotent naïve-hESCs of the inner cell mass (ICM) at the early pre-implantation blastocyst stage and pluripotent formative-hESCs at the peri-implantation blastocyst stage are competent for hPGC induction, while more developmentally advanced pluripotent primed-hESCs of the late post-implantation blastocyst stage are not (Chen et al., 2019; Irie et al., 2015). Only a small subset of conventionally cultured hESCs/hiPSCs maintain a naïve hESC state (Cinkornpumin et el., 2020) while the vast majority remain at the primed hESC state (Messmer et al., 2019), reflecting developmental plasticity between totipotent and primed hESC states (Oosterhuis et al., 2019; Messmer et al, 2019). Given that only a small subset of hiPSCs is capable of hPGCLC differentiation (Irie et al., 2015; Cinkornpumin et al., 2020; Messmer et al., 2019), we tested dose-dependent tumorigenicity by injecting hiPSC_A4_L82 (fig. S1C) at conventional-number (1×10^6^) and high-number (2×10^7^) cell doses into bilateral flanks of three mice per condition. The **3**^**rd**^ **group** used **hEGC-like cells** (**hEGCLCs**) as the surrogate for hEGCs (Kobayashi et al., 2022; Shamblott et al., 1998), which are the in vitro parthenogenic derivatives of unipotent hPGCLCs and hPGCs respectively. Based on normal and teratoma developmental trajectories (Oosterhuis et al., 2019) - and given that VP-MCC is a somatic cancer - we considered the possibility that MST may require parthenogenetic reversion of a MCPyV-transformed hPGC to a pluripotent or totipotent intermediate (Oosterhuis et al, 2019) before somatic neuroendocrine lineage differentiation. This would position hEGCs as a more direct candidate cell of origin than hPGCs, supporting the inclusion of hEGCLCs to test for direct conversion to VMLTs in vivo. It also remained unclear whether MCPyV transfected hEGCLC possess equivalent developmental potency to non-parthenogenetic hPSC counterparts such as MCPyV transfected hiPSCs, particularly in achieving a naïve hESC state to support hPGCLC induction and thus an indirect MST route to VMLTs. To evaluate both possibilities, EGCLC_A4_L82 was injected at both cell doses (1×10^6^ and 2×10^7^) into bilateral flanks of three mice per condition. For both the 2^nd^ group and the 3^rd^ group, MCPyV-negative controls, hiPSC_A4 (provided by the Shioda lab (Kobayashi et al., 2022)) or hEGCLC_A4, were not included as it has been well established that pluripotency alone is coupled to teratoma formation without MST (Damjanov et al., 2016) (Table S1).

## Results

Tumors comprised of both VMLT and teratoma components emerged exclusively from low-number (3×10^5^) hPGCLCs_L82 and from high-number (2×10^7^) hiPSC_L82 injections. Only teratoma component developed in conventional-number (1×10^6^) hiPSC_L82 mice, as well as hECLC_L82 mice regardless of cell numbers (both 1×10^6^ and 2×10^7^). No tumor formation was observed in MCPyV-negative hPGCLC control mice, consistent with their known “unipotency” in vivo from prior studies (Aleckovic et al., 2008; Sugawa et al., 2015). One of the three hPGCLC_A4_L82 mice (#86.1) failed to form any tumor, probably due to a technical failure of cell injection as the mouse was noted to be particularly challenging to apply physical restraint during injection. Bilateral injections of hiPSC_A4_L82 line or hEGCLC_A4_L82 line in the same mouse resulted bilateral mouse xenograft tumors of essentially the same gross and histologic characteristics (Table 1 and Table S1).

**Table 1.**
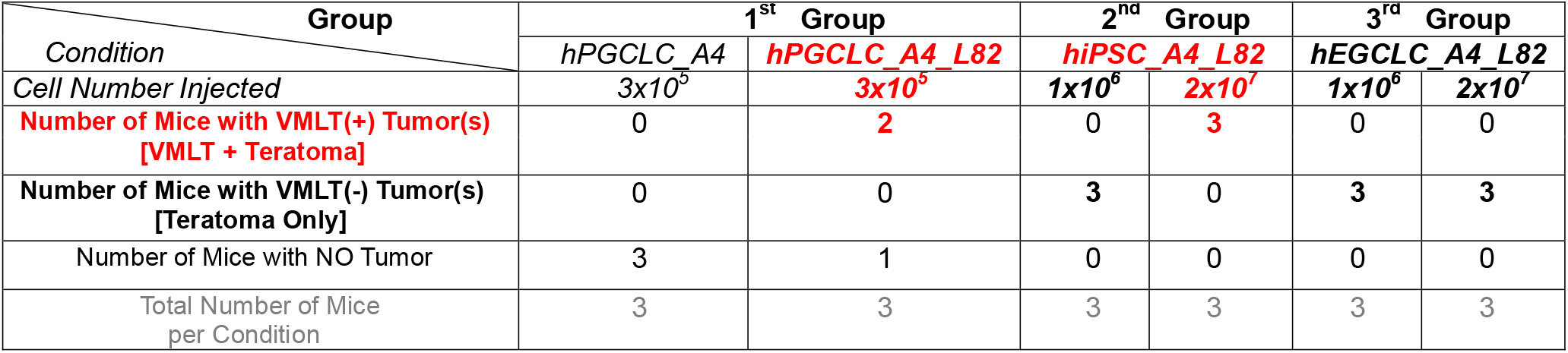
Summary of mouse tumor formation from subcutaneously injected human primeval stem cells.

There was no gross or histological evidence of distant metastasis from any xenograft tumors with VMLTs. This finding is consistent with the known much less aggressive phenotype with only rare distant metastasis from VP-MCC tumor cell line-derived mouse tumors in prior studies (Dresang et al., 2013).

The reason for the marked difference of VP-MCC tumors’ aggressiveness in human versus mouse is unknown and may reflect species differences.

### MST to VMLTs confirmed in VMLT(+) tumors

#### Morphology confirmation of MST to VMLTs

Grossly, VMLT(+) tumors, which were composed of both VMLT and teratoma components, were markedly different from VMLT(-) tumors, which were composed only of a teratoma component. All VMLT(+) tumors were solid and homogeneous with pale and friable texture grossly (Figure 1), while teratoma-only tumors were cystic and mucinous with heterogeneous discoloration (Figure S2). The finding mimicked the typical gross difference between clinical teratomas with MST and pure teratomas (Ulbright et al., 2005). The WHO criteria for MST in human GCTs require a minimum 5 mm expansion of pure somatic type of malignancy component (Berney et al., 2022). Our most salient tumor examples of MST to VMLT were T#85.4, consisting of two discrete tumor nodules with the larger nodule (T#85.4.N1 ~20 mm) composed of pure VMLT tissue (Figure S5A), and T#84.5R ~20 mm, composed almost entirely of VMLT (Figure S6A-B). The rest of the VMLTs also occupied substantial portions of tumors in sizable continuous sheets (~20% - ~80% of tumor masses) (Figure S7-8, Figure S6C, Figure S9A, Figure S20A).

**Figure 1.**
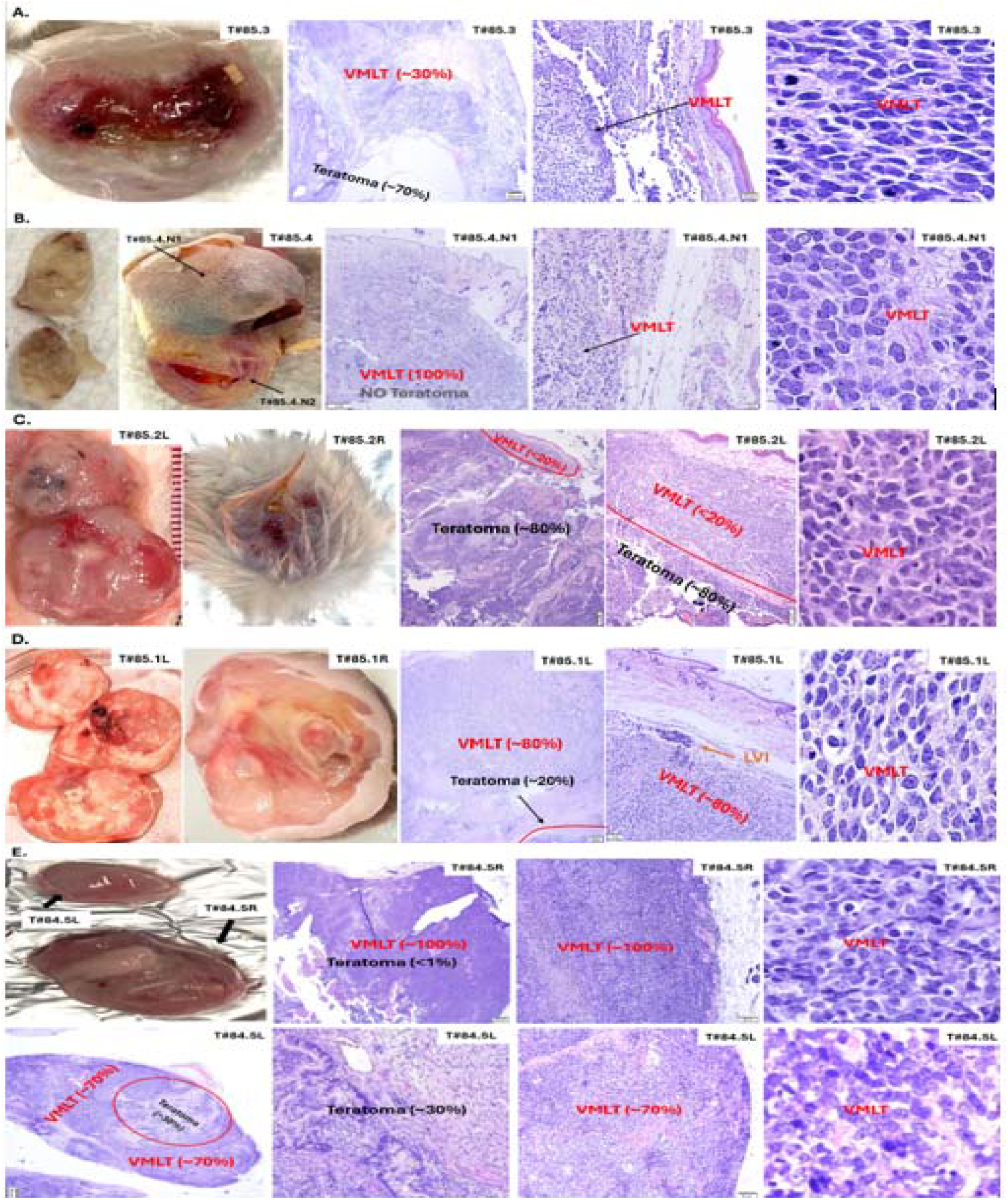
Gross appearance and representative histology of mouse VP-MCC-like tumors (VMLTs) derived from human primeval stem cells: Injected with hPGCLC_A4_L82 (~ 3 × 10^5^ cells): **(A)** Mouse #85.3. **(B)** Mouse #85.4. Injected with hiPSC_A4_L82 (~ 2 × 10^7^ cells): **(C)** Mouse # 85.2. (**D)** Mouse #85.1 (**E)** Mouse #84.5.

Histologically, VMLT(+) tumors exhibited a distinct overall spatial pattern: primitive VMLT components localized superficially and peripherally, surrounding more differentiated teratoma components located deeper and centrally (Figure S6C, S7, S9A, S10A, S18). This organization contrasts with the typical “differentiation gradient” of hPSC-derived mouse xenograft pure teratomas like VMLT(-) tumors, which display a less differentiated core and more mature peripheral regions (Aleckovic et al., 2008). Notably, the teratoma components within VMLT(+) tumors did retain the expected maturation gradient of xenograft pure teratomas (Figure S18A–B). The opposing spatial arrangements of the overall VMLT(+) tumors versus their teratoma components implicate a distinct VMLT tumorigenic pathway requiring proximity to the cutaneous microenvironment.

#### VP-MCC like histopathology of VMLTs

Histologic sections and cytology smears of all VMLTs showed sheets of immature small blue cells with either the typical MCC small round cell cytomorphology in the two hPGCLC_A4_L82 derived tumors T#85.3 or T#85.4 (Figure S5E, Figure S8E, Figure S11), or the atypical MCC large cell variant with admixed larger cells exhibiting prominent nucleoli and more abundant cytoplasm in all six high-number hiPSC_A4_L82 derived tumors, T#84.5L&R, T#85.1L&R, T#85.2L&R (Figure S6F, Figure S9D, Figure S10E, Figure S11). All VMLTs exhibited prominent histological characteristics of malignancy (Figure S12), including extracapsular invasion into dermis, with destruction of dermal striated muscle layer and lymphovascular invasion, with VMLT cells within lymphatic spaces (Figure S7, Figure S8A-D, Figure S6E, Figure S9B-C, Figure S10B-D).

#### VP-MCC like molecular profile of VMLTs

Molecular characterization by RT-qPCR (Figure 2, Table S2) and/or immunohistochemical (IHC) studies (Figure 3, Figure S13, Figure S14-16, Table S3) of VMLTs showed highly similar molecular profiles to each other and to the six VP-MCC cell lines (CVG-1, MKL-1, MKL-2, MS-1, PeTa and WaGa) (Guastafierro et al., 2013; Houben et al., 2013; Houben et al., 2010; Martin et al., 1991; Rosen et al., 1987; Shuda et al., 2008; Velásquez et al., 2018). They were less similar to that of the two virus-negative variant MCC (VN-vMCC) cell lines (MCC-13 and UISO) (Shuda et al., 2008; Ronan et al., 1993; Thomson et al., 1994) that do not express the key neuroendocrine lineage gene INSM1. The VMLTs showed mRNA and/or protein expression in neuroendocrine lineage genes, viral LT antigen, core pluripotency gene and PGC similar to that of VP-MCCs with few exceptions, and a marked departure from that of the corresponding MCPyV transfected primeval stem cells, hPGCLC_A4_L82 or hiPSC_A4_L82 (Figure 2-3, Figure S13, Figure S14-16, Table S2-S3). The high Oct4 expression of all high-number hiPSC_A4_L82-derived VMLTs, in contrast to very low or absent expression in VP-MCC lines (Figure S17), raised concerns about a possible EC component in these VMLTs. However, it was ruled out by PLAP(-) and CD30(-) IHC stains on these VMLTs (Figure S14-16, Table S3). Significantly lower ATOH1 mRNA levels were noted in all VMLTs than VP-MCC cell lines, which might be due to the incipient tumor status of VMLTs versus the advanced cancer stage of these VP-MCC cell lines (Gambichler et al., 2017).

**Figure 2.**
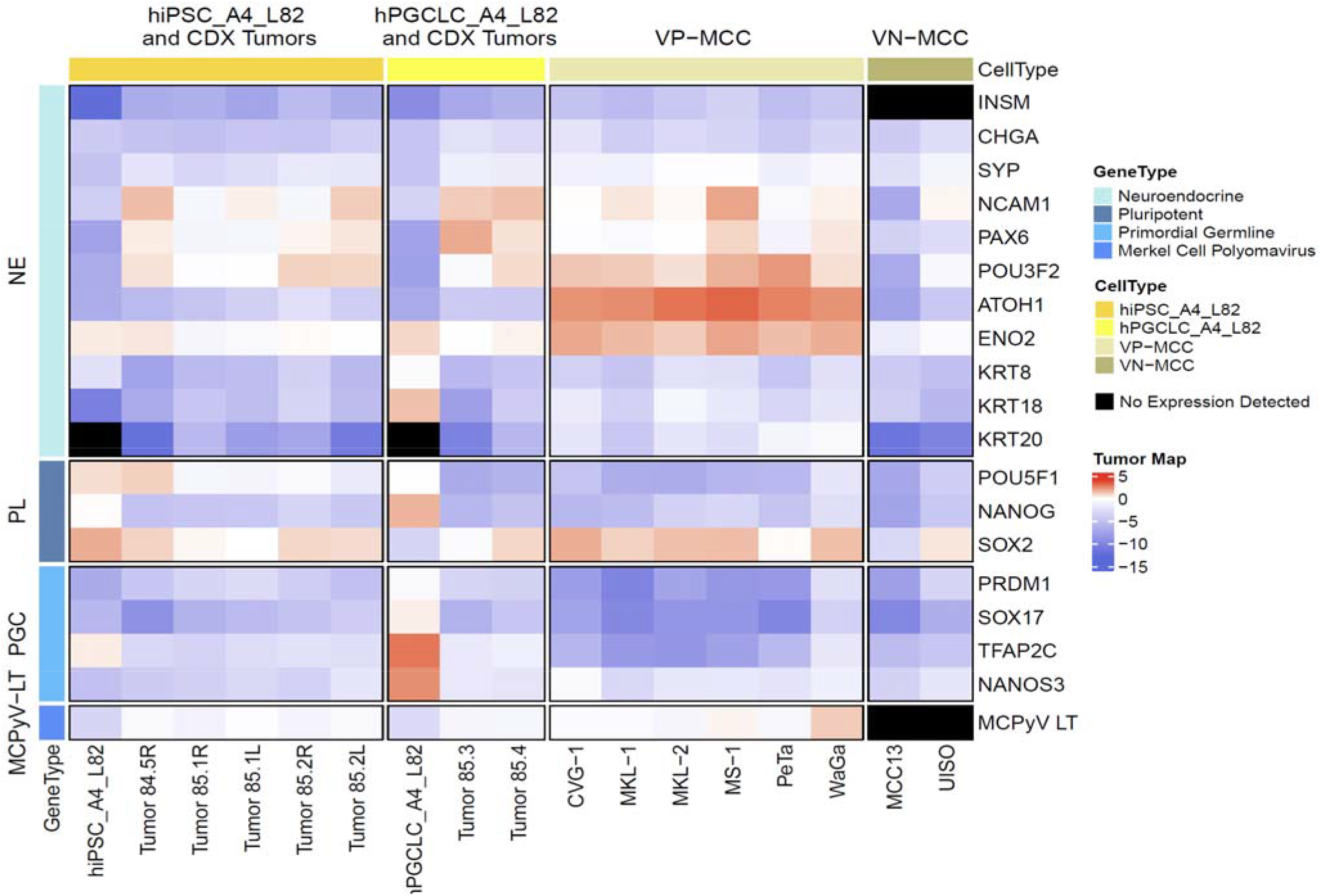
Heatmap of mRNA expression. Comparison of mRNA expression of key neuroendocrine lineage genes, core pluripotent genes, PGC marker genes and McPyv LT viral oncogene among transfected primeval stem cell lines (hiPSC_A4_L82 and hPGCLC_A4_L82), their derivative mouse CDX tumors, VP-MCC and VN-MCC cell lines.

**Figure 3.**
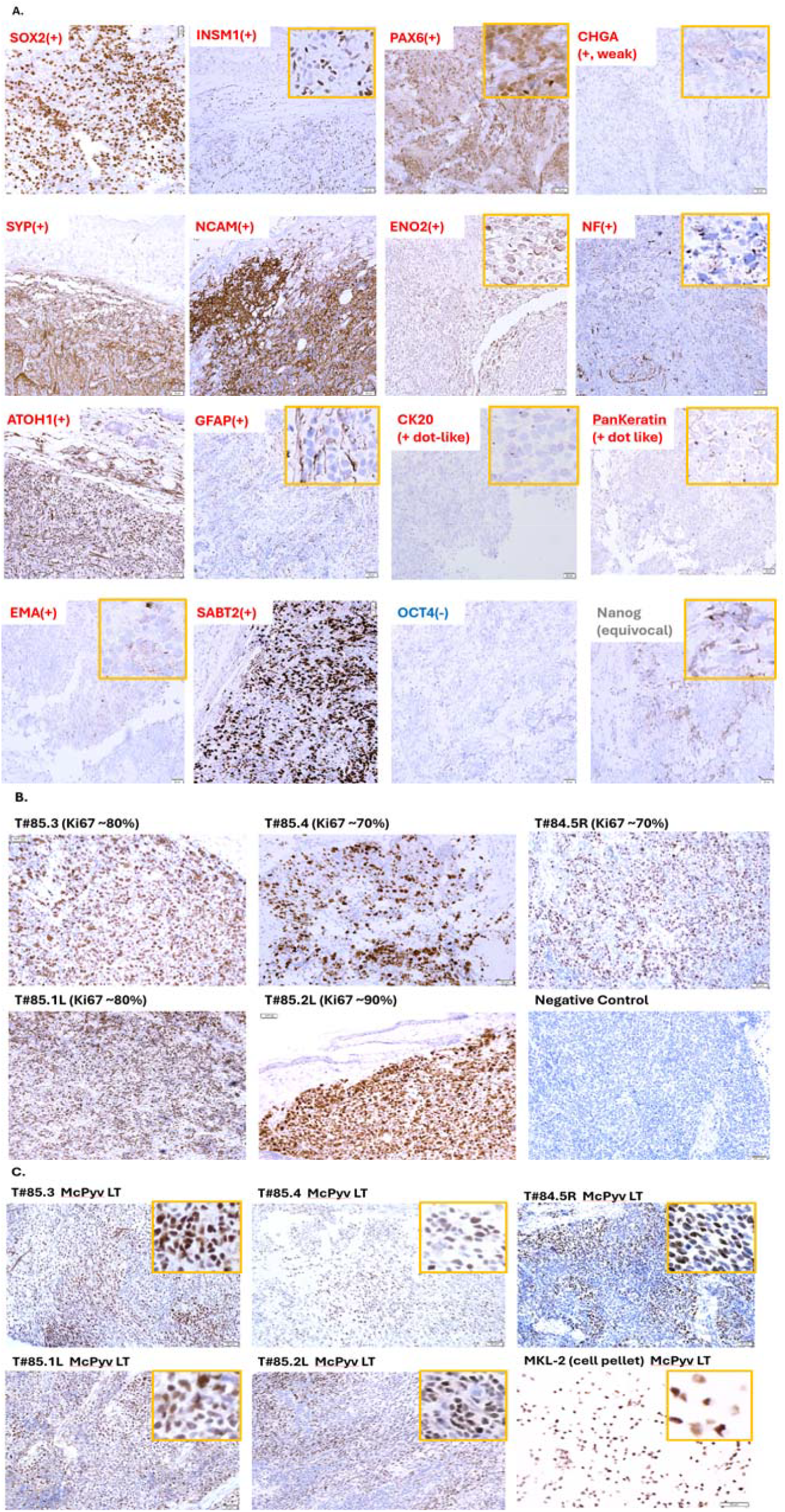
Immunohistochemical studies of mouse VMLTs. **(A)** Protein expression of a panel of genes in VMLT of T#85.4.N1. **(B)** High Ki67 proliferative indexes of VMLTs from five mice. **(C)** Positive expression of McPyv LT oncoprotein in VMLT from five mice. MKL-2 cell pellet was used as a positive control.

#### Global DNA methylation reset before MST

Global DNA methylation (GDM) is dynamic during development, but may serve as a relatively stable epigenetic marker that retains lineage memory. Cancer cells may maintain DNA methylation signatures reflective of their cell of origin despite reprogramming. In normal human development, only two waves of global DNA demethylation occur, each producing a transient hypomethylation nadir during early embryogenesis. The first nadir occurs in totipotent naïve hESC/ICM stage (~40% median GDM); the second nadir - and lowest - in late hPGC stage (~6% median GDM) (Guo et al., 2015; Wen et al., 2019) (Figure S19). The second nadir’s extremely low GDM with near complete GDM erasure is unique to late hPGCs while the first nadir’s moderate DNA hypomethylation characterizes the totipotent naïve hESC/ICM state. In vitro models mirror these developmental states with well-characterized GDM levels: hiPSCs and hEGCLCs (primed hESC-like) show high GDM (~70%), hPGCLCs (early hPGC-like) show intermediate GDM (~50%) (Kobayashi et al., 2022) (Figure S19). GCCs typically reflect this epigenetic hierarchy with few exceptions: seminomas usually match the extremely low GDM of late-hPGCs, ECs usually show moderate global DNA hypomethylation of naïve hESCs, and malignant teratomas usually exhibit higher GDM due to somatic differentiation (Figure 5). All GCCs thus retain variable degrees of global DNA hypomethylation consistent with their late-hPGC origin (Oosterhuis et al., 2019). Somatic cancers are also known to exhibit global DNA hypomethylation, positively correlating with degree of malignancy and cancer progression (Ehrlich, 2009). SCNCs, including MCC, share marked global DNA hypomethylation (Gravemeyer et al., 2022; Harms et al., 2022) with seminomas and ECs - the stem-like compartments of GCCs - as well as stemness and developmental plasticity. To investigate further, we compared GDM in VMLTs, MCC lines, and the only validated seminoma cell line TCam-2 (Mizuno et al., 1993). One important caveat is that DNA methylation is highly sensitive to the microenvironment, and tumor cells maintained in vitro often display artificially elevated GDM (Smiraglia et al., 2001). For instance, the only validated seminoma cell line TCam-2 - despite its composition of transformed late-hPGCs known to exhibit extremely low GDM (~6% median) (Guo et al., 2015; Wen et al., 2019) - shows a moderate GDM level (Nettersheim et al., 2013) comparable to the cell lines of EC mirroring naïve hESC/ICM state^1^ (GDM ~40%)(Guo et al., 2015; Wen et al., 2019), rather than the extreme global hypomethylation observed in primary seminoma tissues (Nettersheim et al., 2013), likely reflecting culture-induced epigenetic drift (Smiraglia et al., 2001).

#### GDM by Flow cytometry and IHC validated

Semi-quantitative flow cytometry and immunohistochemical (IHC) study to measure global 5-methylcytosine (5mC) levels for assessing GDM were validated using reference cell lines with well-characterized GDM profiles. The hiPSC_A4_L82 line, modeling the primed hESC state known to exhibit high somatic-type GDM (~70%) (Kobayashi et al., 2022), showed a very high 5mC signal (MFI ~113K) by flow cytometry. The seminoma cell line TCam-2, which reflects a moderately hypomethylated state similar to ECs or naïve hESCs (GDM ~40%) (Guo et al., 2015; Wen et al., 2019), displayed a much lower 5mC signal (MFI ~7K) by flow cytometry with its isotype negative control yielding a minimal signal (MFI ~3K) and serving as the background reference for absence of GDM (Figure 4A; Table S4). IHC stains of hiPSC_A4_L82 line, TCam2 line and its isotype negative control showed similar trend (Figure 4B1-2,4B6). Positive IHC stains of mouse somatic skin tissue and testis tissue served as positive controls as well (Figure 4B3-4).

**Figure 4.**
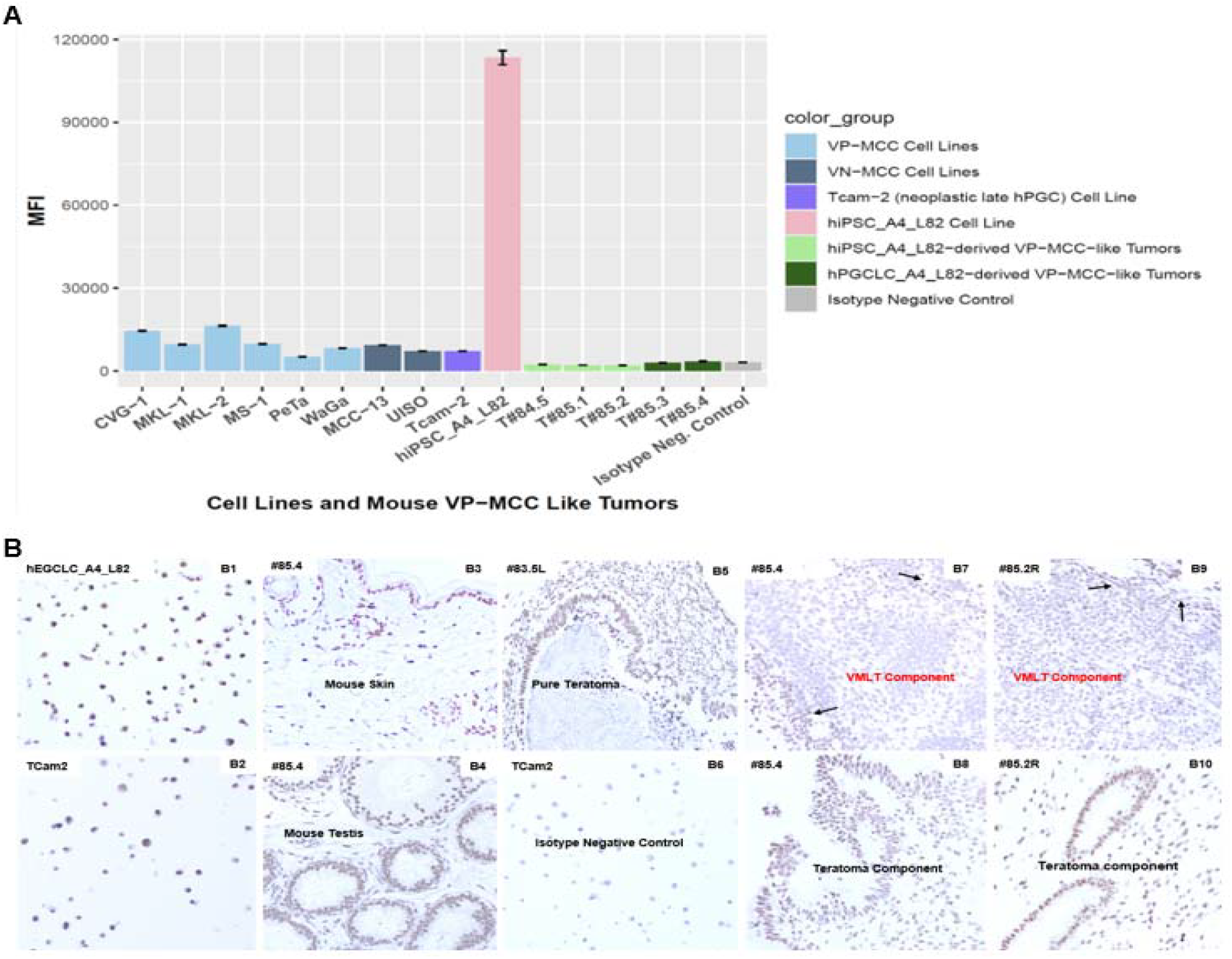
Global 5mC levels to estimate GDM by (A) flow cytometry and (B) immunohistochemical (IHC) studies. **(A)** Flow cytometry estimated global 5mC levels semi-quantitively of hiPSC_A4_L82 cell line, seminoma cell line TCam2 and its isotype negative control, VMLTs from five mice, six VP-MCC cell lines and two VN-MCC cell lines. Global 5mC levels were measured as Median Fluorescence Intensity (MFI) with error bars indicating Standard Error of the Mean (SEM). hiPSC_A4_L82 line and seminoma cell line Tcam2 exhibited very high (~113K) and low (~7K) global 5mC levels indicating GDM levels consistent with previous studies (Kobayashi et al., 2022; Nettersheim et al., 2013). The isotype negative control of TCam2 exhibited extremely low (~3K) global 5mC level as the background signal for absence of GDM, similar to that of VMLTs indicating nearly complete GDM erasure. VP-MCC and VN-MCC cell lines showed low global 5mC levels similar to that of the TCam-2 cell line, which is composed of transformed late-hPGCs. **(B)** Global 5mC IHC studies of hiPSC_A4_L82 cell line (B1), TCam2 line (B2) and its isotype control (B6), as well as the VMLT components from VMLT(+) tumors T#85.4 (B7) and T#85.2R (B9) showed similar findings as that of flow cytometry. Positive IHC stains of mouse#85.4 somatic skin (B3) and testis (B4) served as positive controls.

#### Requisite Late-hPGC state before MST to VMLT

##### VMLTs exhibited unique late-hPGC GDM signature

Flow cytometry analysis of VMLT tumor cells revealed extremely low global 5mC levels (MFI: ~2K–~3.5K) (Figure 4A, Table S4), approaching the isotype negative controls (MFI: ~2K-~3K). Global 5mC IHC studies of VMLT(+) tumors T#85.4 and T#85.2R showed near negative staining for the VMLT components (Figure 4B7, 4B9), and positive staining for the teratoma components (Figure 4B8, 4B10) that is similar to pure teratoma from VMLT(-) tumor T#83.5L (Figure 4B5). These results thus indicated VMLTs exhibited unique late-hPGC GDM signature with near-complete GDM erasure, which suggests a model in which MST occurs directly from the late-hPGC state across the Weismann Barrier (WB), bypassing any intermediary stage marked by moderate to high GDM (Figure S19; Figure 5). The VMLTs were derived either from hPGCLC_A4_L82 (modeling early hPGCs with ~50% GDM) (Kobayashi et al., 2022), or from hiPSC_A4_L82 (modeling primed hESCs with ~70% GDM) (Kobayashi et al. 2022), confirming a profound GDM reset preceding MST to VMLTs.

**Figure 5.**
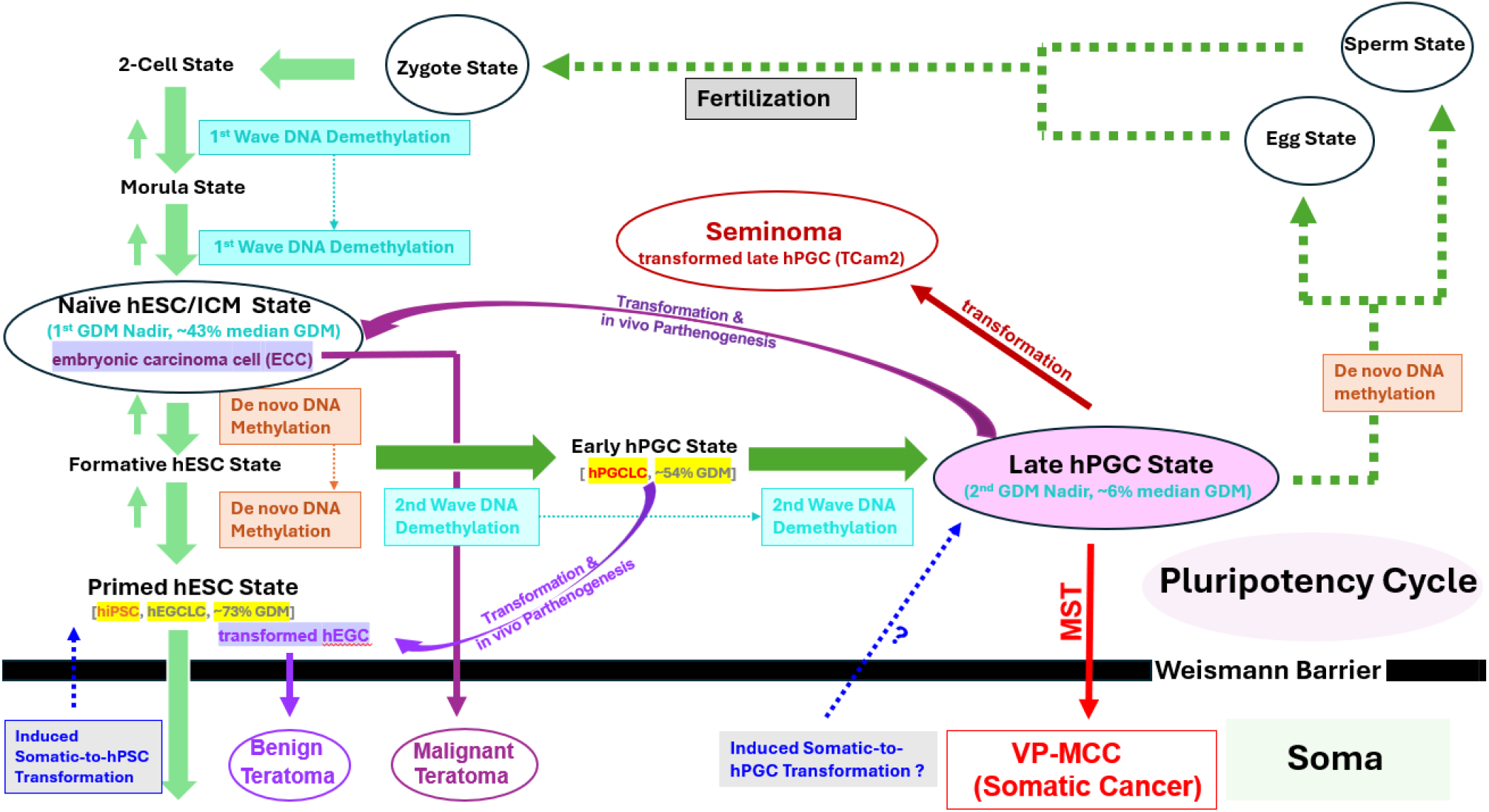
Schematic of normal human embryonic and germline development in comparison to the carcinogenesis of somatic VP-MCC and germ cell tumors (GCTs). Light green arrows show normal early embryogenesis from fertilization to primed pluripotent hESCs before crossing the Weismann Barrier (WB) into somatic differentiation to form the soma. The intrinsic plasticity between the 2-cell stage and the primed hESC stage is reflected in the bidirectionality of the light green arrows (Oosterhuis et al., 2019). Normal human germline development (dark green arrows) begins at the specification of hPGCs from formative hESCs and ends with the formation of sperm or eggs, guided by an intrinsic program suppressing somatic fate, yet exhibiting latent/poised pluripotency (Leitch et al., 2013) with the germline being the only human cell lineage capable of physiological conversion to pluripotency (parthenogenesis) without genetic manipulation. Thus, the germline cycle may be considered a “pluripotency cycle” (Leitch et al., 2013), consisting of both early embryogenesis (light green arrows) and germline development (dark green arrows), which are separated from the soma compartment by the Weismann Barrier (WB). Based on our mouse model, carcinogenesis of somatic VP-MCC (red straight arrow) also crosses the WB to the somatic compartment, similar to teratomas - a type of non-seminomatous GCT (NSGCT) (dark purple and light purple straight arrows) - and normal human embryogenesis (Oosterhuis et al., 2019). However, unlike teratomas or embryogenesis, VP-MCC tumorigenesis crosses the WB directly from the germline (late hPGCs) to the somatic side without a pluripotent intermediary, resulting in mono-lineage (neuroendocrine) somatic differentiation accompanied by developmental arrest. In contrast, tumorigenesis of malignant teratomas more closely mirrors normal embryogenesis, requiring a totipotent EC cell (ECC) intermediate to cross the WB and enable multilineage somatic differentiation. These totipotent stem cells of malignant teratomas are believed to arise via parthenogenetic transformation of transformed hPGCs in vivo (dark purple and light purple curved arrows), rather than from fertilization-derived zygotes. Seminomas, another malignant GCT subtype, consist of transformed late hPGCs and therefore retain the epigenetic signatures characteristic of this stage (Oosterhuis et al., 2019; Wen et al., 2019). Two waves of global DNA demethylation occur during development, each followed by de novo methylation. The first wave begins at the 2-cell stage and reaches its nadir at the naïve hESC/ICM stage, with partial erasure of GDM (~40%) (Guo et al., 2015; Wen et al., 2019). The second, germline-specific wave occurs only during hPGC development, resulting in near-complete GDM erasure and reaching the lowest nadir at the late hPGC stage (GDM ~6%) (Guo et al., 2015; Wen et al., 2019). The moderate global hypomethylation observed in two VN-vMCC cell lines similar to that of seminoma cell line TCam2 and VP-MCC lines suggests a possible somatic-to-hPGC transformation (slanted dashed blue arrow with a question mark), potentially triggered by somatic mutations or other mechanisms. The somatic-to-hPGC transformation may also undergo indirect pathway through reprogramming of somatic cells into hiPSCs followed by specification into a hPGC-like state (vertical dashed blue arrow). The three human primeval stem cell types used in our mouse model - hPGCLCs (modeling early hPGCs), hiPSCs (modeling primed hESCs), and hEGCLCs (modeling hEGCs) - are highlighted in yellow.

##### Direct and indirect tumorigenic pathways to VMLTs

VMLTs’ late-hPGC GDM signature enables delineation of the tumorigenic pathways for VMLTs derived from hPGCLC_A4_L82 versus those derived from high-number hiPSC_A4_L82 cells. A direct pathway, for hPGCLC_A4_L82-derived VMLTs, involved further germline progression from an early-hPGC like state (modeled by hPGCLC_A4_L82) (Kobayashi et al., 2022) to the requisite “late-hPGC” state in vivo, followed by MST to VMLTs (Figure 5). In contrast, an indirect pathway, for hiPSC_A4_L82-derived VMLTs, necessitated an extra initial hPGCLC specification step, in which a small subset of naïve hiPSC_A4_L82 cells first differentiated into hPGCLC_A4_L82 cells. These then proceeded to the “late-hPGC” state before undergoing MST to VMLTs (Figure 5). The observed dosage effect - VMLT formation in high-number (2×10^7^) but not conventional-number (1×10^6^) hiPSC_A4_L82 injections, with no such effect in low-number (3×10^5^) hPGCLC_A4_L82-derived tumors - can be attributed to this additional hPGCLC specification bottleneck. Moreover, clinical observations of rare MST from pure seminomas to somatic malignancies (Dieckmann et al., 2017) including HGNEC intermixed with seminoma cells (Kumaki et al., 2007), offer further clinical support for the plausibility of direct MST from a “late-hPGC”-like state to somatic cancer, reinforcing the pathway proposed here.

#### MCC and seminoma lines show similar GDM

##### VP-MCC and TCam2 lines showed similar GDM

To further validate the findings supporting VP-MCC tumorigenesis via direct transformation from an obligatory “late-hPGC” state, we analyzed global 5mC levels in six VP-MCC cell lines and compared them to the seminoma cell line TCam-2, which consists of transformed late-hPGCs (Oosterhuis et al., 2019).

Flow cytometry showed that VP-MCC lines exhibited moderate global DNA hypomethylation (MFI: ~5K-16K), similar to TCam-2 (~7K), though slightly higher (p<0.027) than VPMLs (~2K-3.5K) (Figure 4A; Table S4). As both VP-MCC lines and TCam-2 were cultured under identical conditions (RPMI + 10% FBS), their mildly elevated 5mC levels relative to that of VMLTs likely reflect epigenetic drift due to in vitro culture conditions as well (Smiraglia et al., 2001).

##### VN-vMCC and TCam2 lines also showed similar GDM

Interestingly, the two VN-vMCC cell lines, MCC-13 and UISO, grown under the same culture condition also displayed comparable levels of moderate hypomethylation (MFI: ~7K-~9K; Figure 4A, Table S4) to that of VP-MCC cell lines, aligning with previous findings (Gravemeyer et al., 2022; Harms et al., 2022). The similarity in GDM between VN-vMCC lines and TCam-2 under matched culture conditions suggests that primary VN-vMCC tumors may also exhibit extreme hypomethylation - an epigenetic hallmark of the “late-hPGC” state observed in primary seminomas. In the case of VN-vMCCs, this “late-hPGC”-like state may arise via reprogramming from somatic cells driven by extensive somatic mutations, including pRB and p53 mutations, characteristic of these tumors (Pedersen et al., 2024; Ma et al., 2023; Harms et al., 2022) (Figure 5).

## Discussion

### Functional evidence against parthenogenic intermediates for MST to VMLT

#### Failure of hEGCLC_A4_L82 to derive VMLT supports no pluripotent intermediate for MST

Failure of hEGCLC_A4_L82 to form VMLTs in vivo indicating a lack of direct conversion to VMLTs from hEGCLC_A4_L82 - the in vitro pluripotent derivative of hPGCLC_A4_L82. This finding therefore provides the functional evidence against the involvement of a parthenogenetic pluripotent hESC-like intermediate in MST to VMLTs, and supports a VMLT tumorigenic pathway that is independent of teratoma formation and uncoupled from parthenogenetic pluripotency.

#### Decreased potency of hEGCLC_A4_L82 suggests no totipotent intermediate for MST

Previous studies showed dichotomous parthenogenic paradigms for normal hPGCs vs transformed hPGCs in NSGCCs (Oosterhuis et al., 2019; Aleckovic et al., 2008; Kobayashi et al., 2022). The developmental potency of parthenogenetic derivatives of normal hPGCs, such as hEGCLCs or hEGCs, has been shown to decrease along the hPGC developmental trajectory: hEGCs derived from post-migratory late-hPGCs lose bona fide pluripotency, whereas hEGCLCs originating from hPGCLCs (modeling early hPGCs) regain pluripotency, as demonstrated by their inability or ability to form teratomas in vivo, respectively (Aleckovic et al., 2008; Kobayashi et al., 2022). Thus, prior entry into the germline and a corresponding decline in developmental potency define the normal hPGC parthenogenesis paradigm. In contrast, transformed hPGCs in NSGCCs exhibit a reversed trajectory, with parthenogenetic derivatives showing increased developmental potency along germline progression: transformed late-stage hPGCs give rise to totipotent EC cells, while transformed early-stage hPGCs yield less potent, primed-state hESC-like derivatives. In this transformed paradigm, prior germline entry is associated with an upregulation of developmental potency - a distinct hallmark of NSGCC transformation (Oosterhuis et al., 2019).

The inability of high-number hEGCLC_A4_L82 cells to form VMLTs in mice, compared to the successful VMLT formation by high-number hiPSC_A4_L82 cells, provides functional evidence that hEGCLC_A4_L82 had failed to transition to the totipotent naïve hESC-like state required for VMLT induction via the indirect pathway through hPGCLC_A4_L82 specification. This reduced developmental potency relative to its non-parthenogenetic counterpart aligns with the normal, rather than NSCGC-transformed, hPGC parthenogenesis paradigm. This observation reflects MCPyV’s inability to functionally enhance parthenogenetic potency in hPGCLCs_A4_L82, in contrast to the developmental potency upregulation characteristic of NSGCC transformation. MCPyV’s apparent incapacity to induce NSGCC-like transformation indicates that hPGCLC_A4_L82 cells, upon viral transduction, cannot parthenogenetically acquire a totipotent EC-like state. These findings are consistent with the GDM evidence supporting the requirement of a “late hPGC” state before direct MST to somatic VMLTs, without the involvement of a totipotent parthenogenetic intermediate. Furthermore, MCPyV’s inability to induce NSGCT-like transformation also suggests that the teratoma components observed in hPGCLC_A4_L82-derived VMLT(+) tumors have arisen directly from an in vivo parthenogenic derivative competent for teratoma formation, from hPGCLC_A4_L82 modeling early-hPGCs (Kobayashi et al., 2022) as spontaneous germline developmental progression in vivo to a “late-hPGC” state would have resulted in a parthenogenic derivative incapable of teratoma formation (Aleckovic et al., 2008).

#### Independent tumorigenic pathways for teratoma vs VMLT

Based on the VMLT tumorigenic pathway supported by the obligatory extremely low GDM unique for “late-hPGC” state and the known intrinsic pluripotency-teratoma coupling with teratoma components exhibiting much higher GDM in VMLT(+) tumors (Figure 4B), an independent tumorigenic pathway is favored for the teratoma components of VMLT(+) tumors, involving multi-lineage somatic differentiation from a pluripotent precursor (Aleckovic et al., 2008). Failure of hEGCLC_A4_L82 to form VMLTs also provided the functional evidence to oppose the involvement of parthenogenic intermediates in VMLT tumorigenesis as discussed above. The independent tumorigenic pathways for the VMLT component versus the teratoma component can provide the rationale for the co-presence of VMLT and teratoma components in this VP-MCC mouse xenograft model, while no mixed VP-MCC and teratoma tumor has been reported in clinical settings. The answers lie in the fact that the experimental mouse teratomas derived from hPSCs are known to be polyclonal (Aleckovic et al., 2008), while clinical cancer is a clonal disease originated from a single transformed cell including VP-MCC that is characterized by a clonal viral insertion site (Feng et al., 2008).

#### Potential tumorigenic functions by MCPyV

The inability of hPGCLC_A4 to form any tumor - teratoma or VMLT - compared to the formation of both components by hPGCLC_A4_L82 implicates several potential tumorigenic functions of MCPyV. These include: (i) promoting survival or anti-apoptotic activity in hPGCLC_A4_L82 (contributing to both VMLTs and teratomas); (ii) enabling parthenogenetic reacquisition of pluripotency (teratomas only); (iii) facilitating progression from an early to late hPGC state (VMLTs only); and (iv) directly transforming late-hPGC-like cells into VMLTs (VMLTs only). Given MCPyV’s inability to induce NSGCT-like transformation and the established teratoma-forming capacity of normal hEGCLCs, its role in teratoma formation is likely minimal. Rather, as extragonadal hPGCs are capable of spontaneous germline differentiation into late-hPGCs or beyond in non-gonadal environments before undergoing apoptosis (Oosterhuis et al., 2019), the most plausible MCPyV-driven mechanisms are enhanced survival and direct transformation of late-hPGC-like cells into VMLTs. One mechanism for MCPyV driving MST from late-hPGCs to VMLTs could be through its inhibition of p53 indirectly (Pedersen et al., 2024) as inactivating mutations of p53 have been associated with MST (Oosterhuis et al., 2019). Further studies are warranted to elucidate the molecular mechanisms underlying MCPyV-mediated VMLT formation.

#### Early developmental vs carcinogenic pathways

Notably, the “late-hPGC” state is also a requisite intermediate in GCCs, indicating overlapping cancer biology between GCCs and VP-MCC, a somatic cancer. This convergence is exemplified by the globally hypomethylated DNA epigenetic profile - a recognized molecular hallmark of cancer observed across GCCs and somatic cancer alike (Oosterhuis et al., 2019; Ehrlich, 2009). On the other hand, our model supports a fundamental divergence between normal human development and VP-MCC carcinogenesis at the level of the Weismann Barrier (WB). In canonical early human development, pluripotent hESCs generate somatic tissues through orderly, multi-lineage differentiation. In contrast, VP-MCC appears to originate from a “unipotent” late-hPGC with known intrinsic germline suppression of somatic fate, yet bearing latent or poised pluripotency (Leitch et al., 2013), that aberrantly crosses the WB into the somatic compartment, differentiating exclusively along a neuroendocrine lineage and undergoing developmental arrest. These findings suggest that VP-MCC may arise via an aberrant ontogenic process in which the germline directly gives rise to a somatic malignancy - akin to the emergence of a highly abnormal new life form (Figure 5). This MST model also supports key distinctions between somatic cancers such as VP-MCC and GCCs exhibiting multilineage somatic differentiation, namely malignant teratomas. The latter more closely parallels normal human embryogenesis, requiring a totipotent embryonal carcinoma (EC) cell intermediate to traverse the WB and undergo multilineage somatic differentiation with aberrant developmental arrest. These EC cells are thought to arise via parthenogenetic transformation of late-stage hPGCs in vivo, rather than from fertilization-derived zygotes. Future investigations are needed to determine whether this unifying germ cell theory-based paradigm applies more broadly to other somatic malignancies. For instance, in VN-vMCCs, somatic mutations may induce a functionally equivalent “late-hPGC” state, substituting for the native “late-hPGC” state observed in VP-MCCs.

## Conclusion

In summary, we developed an MST mouse model demonstrating MCPyV-driven transformation of hPGCLCs into somatic VMLTs. GDM analyses identified a “late-hPGC” state as a necessary developmental precursor for direct MST to VMLTs. Spatial, epigenetic, and functional data collectively support the existence of independent tumorigenic trajectories for the VMLT and teratoma components within the same VMLT(+) tumor, and oppose a parthenogenetic pluripotent or totipotent intermediate in VMLT tumorigenesis. This model thereby establishes a direct developmental link between late-hPGC and a somatic cancer, supporting the same cell (or cell state) of origin for both GCCs and somatic cancer.

Finally, the genetic simplicity and experimental tractability of our VP-MCC–based MST model provide a robust platform for elucidating the molecular mechanisms driving VP-MCC pathogenesis. This system may also yield critical insights into the molecular underpinnings of VN-MCC and other treatment-refractory high-grade neuroendocrine carcinomas (HGNECs) characterized by high mutational burdens and genetic complexity. By directly establishing a developmental origin for VP-MCC, this model supports a shift toward a developmental biology-driven framework for advancing somatic cancer research beyond the mutation-centric and soma-centric paradigm.

Positive IHC stains are observed in the teratoma components of VMLT(+) tumors T#85.4 (B8) and T#85.2R (B10), similar to that of pure teratoma of VMLT(-) tumor T#83.5L (B5). Scattered positive cells (black arrows), next to the VMLT tumor cells exhibiting near negative IHC staining, in VMLT(+) T#85.4 and T85.2R tumors are mouse somatic white blood cells (B7) or dermal stromal cells (B9) respectively.

## Method

### Cell Culture

The human iPSC A4 line (hiPSC_A4) generated from commercially obtained Caucasian male (46XY) neonatal foreskin fibroblasts and the derived GFP+ long-term-culture human primordial germ cell like cell (LTC-hPGCLC) A4 line (hPGCLC_A4) were gifted by the Shioda lab (Kobayashi et al., 2022). The human embryonic germ cell like cell A4 line (hEGCLC_A4) were derived from hPGCLC_A4 based a 10-day conversion protocol facilitated by SCF and FGF2 from the Shioda lab (Kobayashi et al., 2022). Both hiPSC and hEGCLC cultures were maintained in the Essential 8 medium (Gibco, Cat#A1517001) or occasionally the Essential 8 Flex medium (Gibco, Cat#A2858501) on human ESC-qualified Matrigel (Corning, Cat#354277) and passaged at every 3-7 days using ReLeSR (StemCell Technologies, Cat#100-0483) in the presence of the ROCK inhibitor Y-27632 (StemCell Technologies, Cat#72304).

Karyotyping analysis were performed every 10-15 passages to assure the genetic integrity of hiPSCs, hEGCLCs and hPGCLCs and their L82 transfected counterparts (Figure S20). The six VP-MCC cell lines, CVG-1, MKL-1, MKL-2, MS-1, PeTa and WaGa were cultured in RPMI 1640 medium (Gibco, Cat#11875093) with 10% FBS (Sigma, Cat#F2442-500ML) as suspension cultures, and passaged every 3-4 days by 1:2 to 1:10 split. The two VN-vMCC cell lines MCC-13 and UISO as well as the TCam2 cell line were also cultured in RPMI 1640 medium (Gibco, Cat#11875093) with 10% FBS (Sigma, Cat#F2442-500ML) as adherent cultures, and passage every 3-4 days by incubation with 0.25% trypsin from Trypsin (Gibco, Cat#15090046) for up to 8 minutes. Mycoplasma contamination testing was performed every six months or when there was a suspicion for contamination was ruled out by performing a PCR based mycoplasma detection kit based on manufacturer’s protocol (PCR mycoplasma detection kit (Thermo Scientific, Cat#J66117.AMJ).

### Derivation of MCPyv+ primeval stem cell lines hPGCLC_A4_L82, hiPSC_A4_L82 hEGCLC_A4_L82

The lentivirus virus vector plenti.puro.MCV.ER.RAZ.2 (a gift from Reety Arora & Sudhir Krishna Addgene plasmid #114382; http://n2t.net/addgene:114382; RRID:Addgene_114382) abbreviated as “L82” containing a C-terminus truncated MCPyv genome and puromycin resistance gene, a 2^nd^ generation packaging vector psPAX2 (AddGene#12260) and an envelope vector pMD2.G (AddGene#12259) were purchased from AddGene.Org. Lentivirus productions of MCPyv containing lentivirus were done with FuGene 6 transfection reagent (Promega, Cat#E2691) and OptiMem serum free medium (Invitrogen Cat#31985). Stable lentivirus transfections followed by puromycin selection to produce MCPyv integrated derivatives hPGCLC_A4_L82 and hiPSC_A4_L82 followed standard AddGene protocol with polybrene (Santa Cruz Biotechnology, Cat#sc-134220) and minor modification due to the requirement of daily Essential 8 medium change for pluripotent stem cell line hiPSC_A4. MCPyv integrated hEGCLC_A4_L82 was derived from hPGCLC_A4_L82 following the same 10-day in vitro conversion protocol (Kobayashi et al., 2022) facilitated by SCF and FGF2 used for converting hPGCLCs to hEGCLCs.

### MCPyv+ human primeval stem cell line-derived mouse xenograft study

All animals used in this study were handled in accordance with an animal study protocol approved by the Uniformed Services University of Health Sciences Institutional Animal Care and Use Committee. As MCC occur in adults with a male dominance, 4–6-week-old adult and male immunocompromised NSG mice [NOD.Cg PrkdcscidIl2rgtm1Wjl/SzJ, Jackson Laboratory Stock# 005557] were used for the study. Simple physical restraint with double-hand method was used for injecting cells subcutaneously to mouse flank.

Mice were monitored regularly for health status and were euthanized by CO2 asphyxiation upon reaching humane endpoint criteria. Three mice were used for testing tumorigenicity for each group of total six groups. For the first two groups, only 3×10^5^ cells/injection of the hPGCLC_A4 line or the hPGCLC_A4_L82 line was injected to the left flank of three mice for each cell line. For the hiPSC_A4_L82 line, 1×10^6^ cells/injection and 2×10^7^ cells/injection were injected to bilateral flanks of three mice in two groups. The same injections were also performed for hEGCLC_A4_L82 in two groups as for hiPSC_A4_L82. All human primeval stem cells were suspended in 50 ul with 10 μM ROCK inhibitor Y-27632 and mixed with an equal volume of Matrigel (Corning, Cat#354277) before injected subcutaneously into the flanks of the mice (Prokhorova et al., 2009; Kobayashi et al., 2022). Mice were monitored regularly for tumor development and sacrificed upon reaching humane endpoint criteria.

Tumors were collected by dissection; friable portions of tumors were collected in sterile RPMI to prepare cell suspension by mechanical disruption using scissors and forceps. Spun down cell pellets by centrifuge at 400xg were then used for both single cells suspension for flow cytometry and to extract total RNA using the PureLink RNA extraction mini kit (Thermo Fisher, Cat#2592159), as well as cryopreserved in FBS (Sigma, Cat#F2442-500ML) with 8% DMSO (Thermo Fisher, Cat#M3148-250ml). Cryopreserved tumor cells were later thawed to extract DNA by PureLink™ Genomic DNA Mini Kit (Thermo Fisher, Cat#182001). The rest of the tumors were fixed 4% paraformaldehyde, and subjected to the standard H&E as well as IHC staining. Cell line authenticity for all VMLT (+) tumors was confirmed by no variation at any loci of five mononucleotide repeats and two pentanucleotide repeats (NR-21, BAT-26, BAT-25, NR-24, MONO-27; and Penta C, Penta D) among themselves and all three human primeval stem cell lines (hPGCLC_A4_L82 and hiPSC_A4_L82), using Promega OncoMate MSI Analysis System (Cat. # MD2140; Madison, WI) and DNA extracted from the tumor cells and cell lines (Figure S21).

### RT-qPCR

Total RNA of cell lines was also extracted using the PureLink RNA extraction mini kit (Thermo Fisher, 2592159). Reverse transcription was performed using Luna one-step RNA-to-Ct RT-qPCR kit (New England Biolab, E3005E), 5 uM gene-specific primer pairs and 5ng RNA per final 20-ul RT-qPCR reaction and Azure Ciel 3 qPCR detection system (Azure Biosystems, CA) with the cycling/melting curve RT-qPCR program recommended by the manufacturers. All primer sequences are listed in table S4. mRNA expression of each gene was normalized relative to the mRNA expression of housekeeping gene RnaseP as ΔCq. All RT-qPCR experiments were repeated at least three times.

### Immunohistochemistry (IHC) Study

Sections of CDX mouse tumors or pellets of cell lines were first deparaffinized by incubating in xylene, followed by rehydration with ethanol solutions (100% and 95%) and distilled water. Antigen retrieval was performed using EDTA Antigen Retrieval Buffer (pH 8.0, 10X) (Vitro Vivo Biotech, Cat#VB-6007) in a steamer, with sections boiled for 40 minutes-1 hour in 1X EDTA Buffer (Table S6). After washing, endogenous peroxidase activity was blocked with 3% hydrogen peroxide prepared from Hydrogen Peroxide 30% Solution (Lab Alley, Cat#HPL30) and sections were then blocked with 5% goat serum (Gibco, Cat#16210064) in TBST (QuickSilver Powdered Buffer Packs, Cat#EB-1203) for 1 hour. Primary antibodies were diluted based on manufacturer’s recommendation and empirical testing (table S5) and the Rabbit IgG isotype control (Invitrogen, Cat# 31235) or mouse IgG isotype control (Invitrogen, Cat#31903) was diluted to the same final concentrations. If the primary antibodies are of rabbit host, they were incubated overnight at 4°C with the only exception of 5mC antibody (Invitrogen, Cat#MA5-24694) for one hour at room temperature. Following antibody incubation, SignalStain® Boost IHC Detection Reagent (HRP, Rabbit) (Cell Signaling Technology, Cat#8114S) was applied per the manufacturer’s instruction, and sections were incubated for 30 minutes. However, if the primary antibodies were of mouse host, to exclude false positive signal from mouse tissue, the sections were stained for primary and secondary antibodies by M.O.M.® (Mouse on Mouse) ImmPRESS® HRP (Peroxidase) Polymer Kit (Vector Laboratories, Cat#MP-2400) per the manufacturer’s protocol.

SignalStain® DAB Substrate Kit (Cell Signaling Technology, Cat#8059S) was applied to visualize the antigen-antibody complex, and counterstaining was performed with Hematoxylin (Cell Signaling Technology, Cat#14166S). The sections were subsequently dehydrated through 95% and 100% ethanol followed by xylene before being mounted with SignalStain® Mounting Medium II (Cell Signaling Technology, Cat#84583S) under a coverslip. Semi-quantitative 5mC IHC stains were performed three times on cell lines, tumor sections, mouse somatic tissue positive controls and isotype negative controls.

### Flow cytometry 5mC Analysis

Single cell suspensions were prepared for VMLT+ mouse tumors, VP-MCC cell lines and hiPSC_A4_L82 line, as well as VN-MCC cell lines and TCam2 cell line via different methods. Single cell suspensions were made for friable mouse VMLTs by mechanical disruption with scissors and forceps in sterile RPMI. Single cell suspensions of floating VP-MCC cell lines and attached hiPSC_A4_L82 line required cell separation by incubation with ReLeSR™ (Stem Cell Technologies, Cat#100-0483) for 20+ minutes and up to 10 minutes at 37°C respectively. Single cell suspensions of attached VN-MCC cell lines and TCam2 cell line required cell separation by incubation with 1.25% trypsin (Gibco, Cat#15090046) for up to 8 minutes. All separated cells were then filtered through 100 um cell strainers (Ibis Scientific, Cat#229486). Prepared single cell suspensions were then stained with eBioscience™ Fixable Viability Dye eFluor™ 780 (Invitrogen, Cat#65-0865-14) for 30 minutes in a blocking buffer of 2% FBS (MilliporeSigma, Cat#F2442-500ML) in PBS (Cytiva, Cat#SH30256.02). eBioscience™ Foxp3/Transcription Factor Staining Buffer Set (Invitrogen, Cat#00-5523-00) was used for cell fixation and antibody staining per the manufacturer’s protocol. Briefly, the cell suspensions were fixed and stained with 5-Methylcytosine (5mC) Recombinant Rabbit Monoclonal Antibody (RM231) (Invitrogen, Cat#MA5-24694) at 1:1000 final dilution and/or the Rabbit IgG Isotype Control (Invitrogen, Cat#31235) at the same final concentration for 30 minutes on ice, followed by staining with Goat anti-Rabbit IgG (H+L) Highly Cross-Adsorbed Secondary Antibody, Alexa Fluor™ 790 (Invitrogen, Cat#A11369) at 1:500 final dilution for 30 minutes. The cells were then stained with Alexa Fluor™ 594 conjugated Rabbit Sox2 antibody (Santa Cruz Biotechnology, Cat# sc-365823) for 30 minutes on ice. Cell suspensions were then placed in the eBioscience™ Flow Cytometry Staining Buffer (Invitrogen, Cat#00-4222-26) and analyzed by flow cytometer Cytek Aurora.

FlowJo software was used to analysis flow data by gating the Sox2(+) population for VMLT cells and the Median Fluorescence Intensity (MFI) at Alexa Fluor™ 790 as well as the Standard Error of the Mean (SEM) were calculated by the FlowJo.

### Statistical Analysis

All RT-qPCR experiments were repeated at least three times. p values were calculated by either paired or unpaired one tail Student T-test. A p-value with <0.05 was considered statistically significant. Based on a binomial distribution of single injection success rate of 80-100% for teratoma formation from subcutaneous injection of hPSCs mixed with Matrigel (Prokhorova at el., 2009), we estimated three-mice per group being the minimum number to ensure the achievement of a high success rate (99.2%) of tumor formation.

## Supporting information

Supplemental without Table S2

## Resource availability

### Lead contact

Any correspondence and material requests should be addressed to corresponding author Wendy Yang.

### Material availability

hPGCLC_A4 and hiPSC_A4 cell lines and their derivative cell lines are subject to the restrictions stated in a materials transfer agreement (MTA) with Dr. Toshi Shioda lab at Massachusetts General Hospital.

### Data and code availability

All data are available in the main text or the supplementary materials.

## Acknowledgement

We would like to thank Toshi Shioda for providing LTC-hPGCLC_A4 and hiPSC_A4 cell lines, technical guidance and protocols; Yuan Chang, Patrick Moore, Masahiro Shuda, Isaac Brownell and James DeCaprio for providing virus-positive and virus-negative Merkel cell carcinoma cell lines, technical guidance, and/or protocols; Alan Epstein for providing TCam2 cell line and protocols; Reety Arora and Sudhir Krishna for AddGene plasmids; Joel Moncur for mouse xenograft tumor histopathological evaluation and manuscript review; Guanghua Wang for cell line and mouse xenograft tumor authentication, mouse tumor histopathological and manuscript review; Geeta Upadhyay for technical advices and manuscript review; Steve Dumler and Dennis Grab for manuscript review; as well as the USUHS Genomics Core and Flow Cytometry Core Lab in Biomedical Instrumentation Center for DNA primer synthesis and flow cytometry services respectively. This study was funded by Uniformed Services University of the Health Sciences StartUp fund.

## Author Contributions

Conceptualization: WY; Methodology: WY; Investigation: WY, SR; Visualization: SC, WY; Funding acquisition: WY; Project administration: WY; Supervision: WY; Writing - original draft: WY; Writing - review & editing: WY, SC

## Ethics Declaration

The authors declare no competing interests.

## Disclaimer

The opinions and assertions expressed herein are those of the author(s) and do not reflect the official policy or position of the Uniformed Services University of the Health Sciences or the Department of Defense.

## Supplementary Information

Figures S1 to S21

Tables S1 to S6

During the preparation of this work the author(s) used ChatGPT in order to improve language and readability. After using this tool/service, the author(s) reviewed and edited the content as needed and take(s) full responsibility for the content of the publication.

