## Supplemental without Table S2 for "A polyomavirus-positive Merkel cell carcinoma mouse model supports a unified cancer origin"

**The supplemental information includes:**

Figures. S1 to S21

Tables S1, S3, S4, S5, S6

**Other Supplementary information for this manuscript include the following:**

Table S2 (in Excel file)

Figures S1-S21

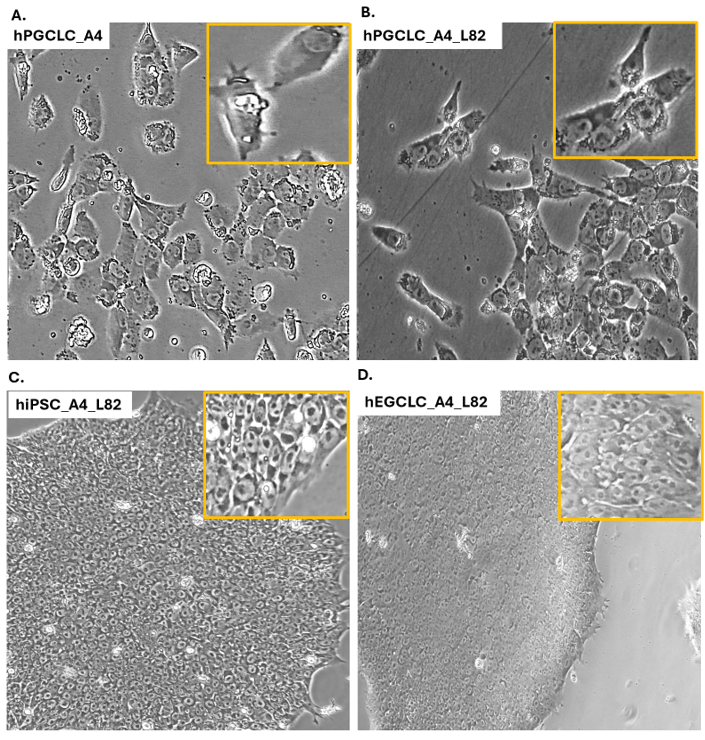

**Figure S1.** **Cell cultures of four primeval stem cell lines injected into NSG mice.**

**(A)** hPGCLC_A4 **(B)** hPGCLC_A4_L82 **(C)** hiPSC_A4_L82 **(D)** hEGCLC_A4_L82

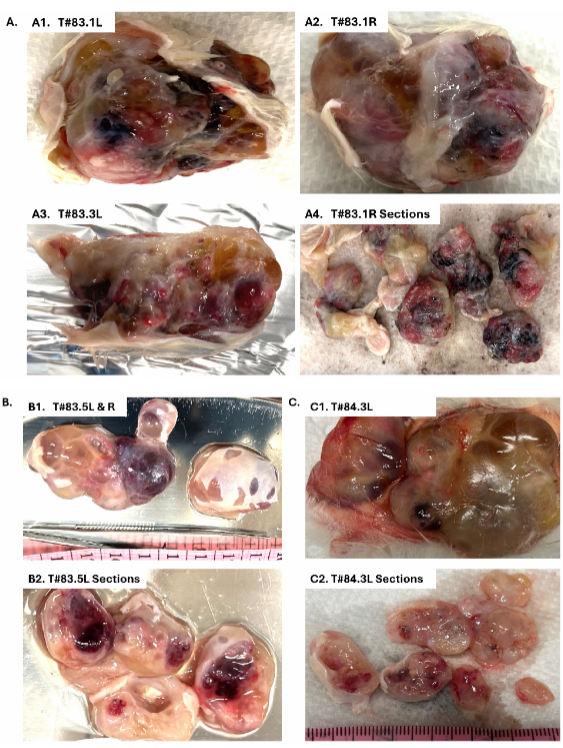

Figure S2. Gross appearance of mouse tumors composed of teratoma only. Tumors induced by

injection of (A) conventional number of hiPSC-A4_L82 cells: T#83.1L, T#83.1R and T#83.3L

(B) conventional number of hEGCLC_A4_L82 cells: T#83.5L (C) very high number of

hEGCLC_A4_L82 cells: T#84.3L

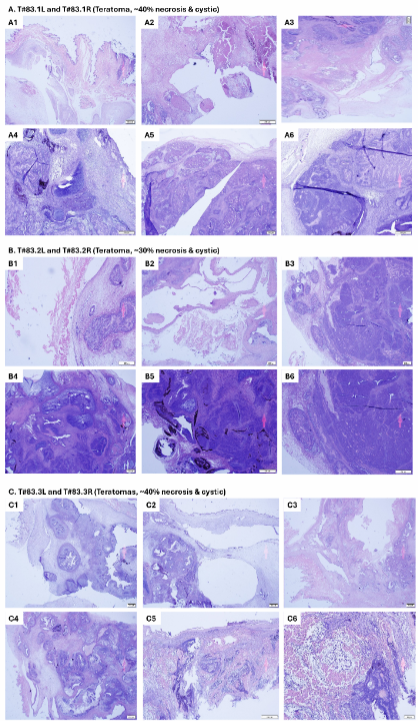

Figure S3. Bilateral teratomas formed after hiPSC_A4_L82 injection. Mice were injected with conventional

numbers (1×10 6 cells) of hiPSC_A4_L82. (A) mouse #83.1 (B) mouse #83.2 (C) mouse #83.3. Frequent

foci of cyst formation and necrosis are noted. Although some areas of immature neuroectodermal tissues

were noted, no large infiltrating sheets of small blue cells to suggest malignant somatic transformation to

MCC-like tumors were identified. The teratomas exhibited no malignant histological features with round

and lobulated rather than infiltrating borders.

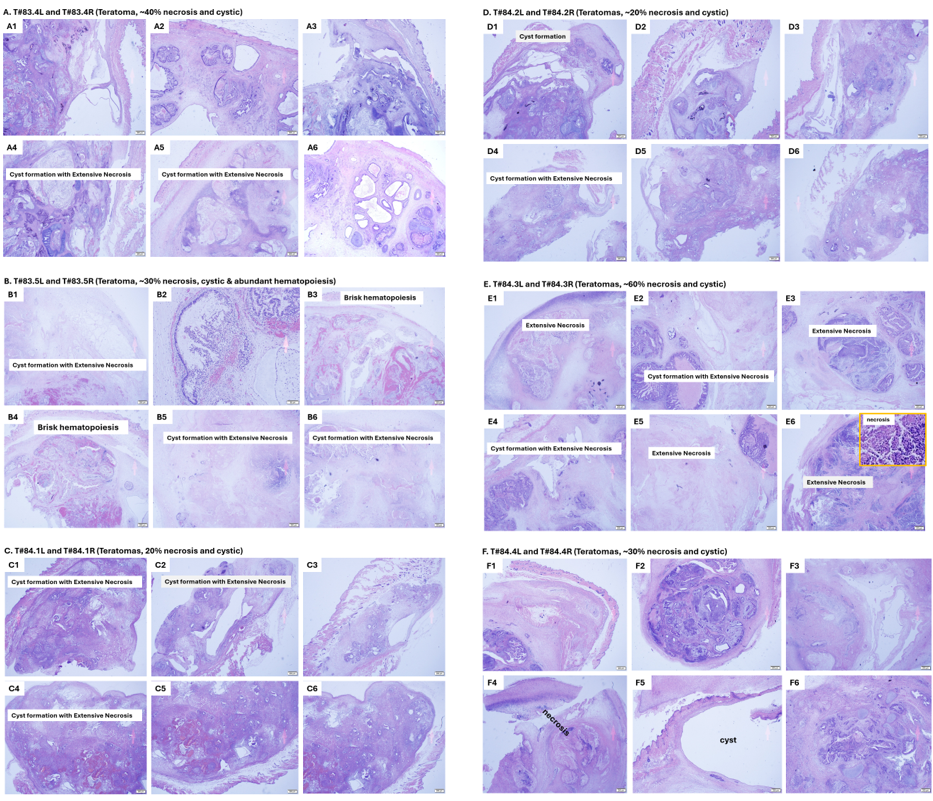

**Figure S4. Bilateral teratomas formed** **after hEGCLC_A4_L82 injection.** Mice were injected with conventional numbers (1×10^6^ cells) of hEGCLC_A4_L82 **(A)** mouse #83.4 **(B)** mouse #83.5 **(C)** mouse #84.1, or with very high numbers (2×10^7^ cells) of hEGCLC_A4_L82 **(D)** mouse #84.2 **(E)** mouse #84.3 **(F)** mouse #84.4. All hEGCLC_A4_L82 derived tumors, despite the difference in numbers of cells injected, showed similar histology and resembled the teratomas derived from injections with conventional numbers of hiPSC_A4_L82, with frequent cyst formation, foci of necrosis and no evidence of malignant somatic transformation to MCC-like tumors. Brisk hematopoiesis was noted in bilateral teratomas of mouse #83.5.

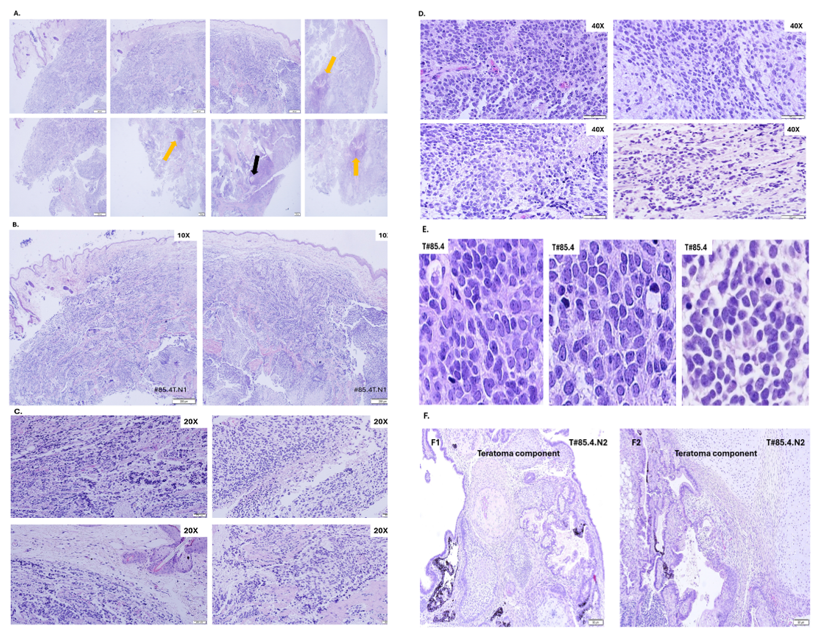

**Figure S5.** **Representative sections of T#85.4.N1 & T#85.4.N2** **(A)** Serial sections of the entire T#85.4.N1 at 2X consisting of pure VMLT in sheets or occasional trabecular growth pattern (orange arrows) with a single tiny focus of epidermal squamous differentiation (black arrow) and no teratoma component. **(B)** Representative section of T#85.4.N1 at 10X. **(C)** Representative section of T#85.4.N1 at 20X. **(D)** Representative section of T#85.4.N1 at 40X. **(E)** High power fields (HPFs) show VMLT cells in both T#85.4.N1 and T#85.4.N2 with typical cytologic features of MCC including high N/C ratio, fine and salt pepper chromatin, inconspicuous nucleoli and frequent mitotic figures. **(F)** Teratoma components of T#85.4.N2 with ectodermal (squamous epidermal), mesodermal (cartilage) and endodermal (glands) differentiation.

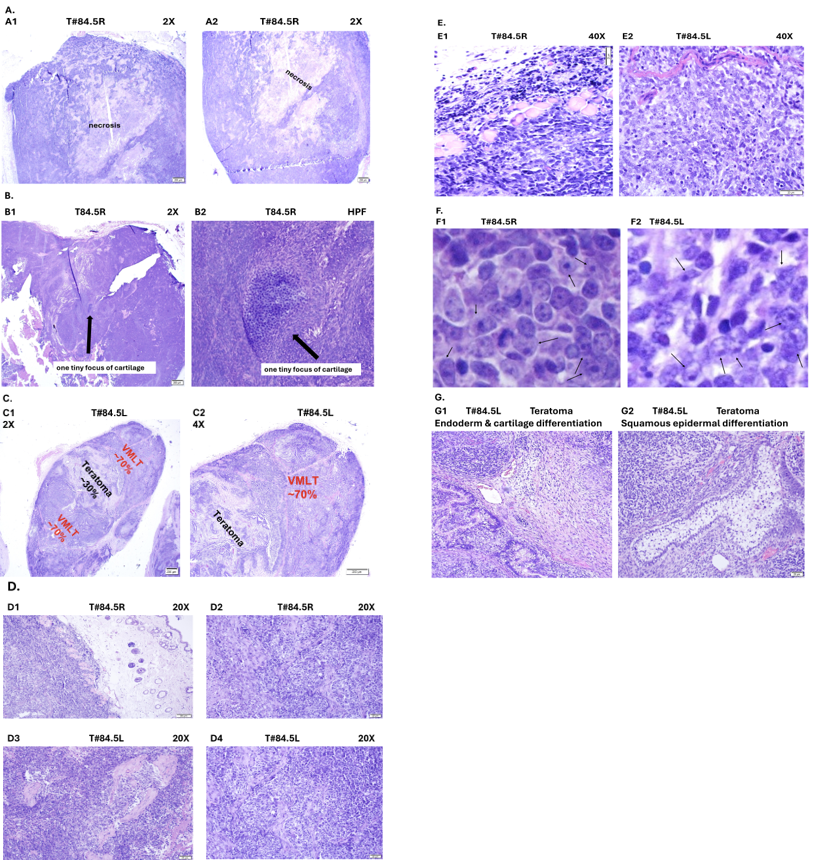

**Figure S6.** **Sections of VMLTs of T#84.5R and T#84.5L.** **(A)** Two representative sections covering the entire T#85.1R tumor show only VMLT with central necrosis and no teratoma component (2X). **(B) (B1)** Representative section of the entire T#84.5R tumor with all VMLT except for a tiny focus of cartilage in the center (black arrow) (2X). (**B2)** HPF shows detail of the tiny single focus of cartilage. **(C)** Representative section of the entire T#84.5L tumor shows a minor central teratoma component (~30% of tumor) surrounded by an extensive peripheral VMLT component (~70% of the tumor). **(D)** Sections of VMLT in T#84.5R and T#84.5L show sheets of small blue cells, with dermal invasion shown in **D1** (20X). **(E)** Sections of VMLT of T#84.5R and T#84.5L show VMLT cells infiltrated through mouse dermal striated muscle layer when invaded into dermis (40X). **(F)** HPF views of cytomorphology of VMLT cells in T#84.5R **(F1)** and T#85.1L **(F2).** As well as typical VP-MCC-like cells with salt and pepper chromatin and inconspicuous nucleoli, intermixed larger atypical cells with paler vesicular chromatin, prominent large nucleoli and more abundant cytoplasm are identified (black arrows). **(G)** Sections of central teratoma component with three germ layer differentiation in T#84.5L.

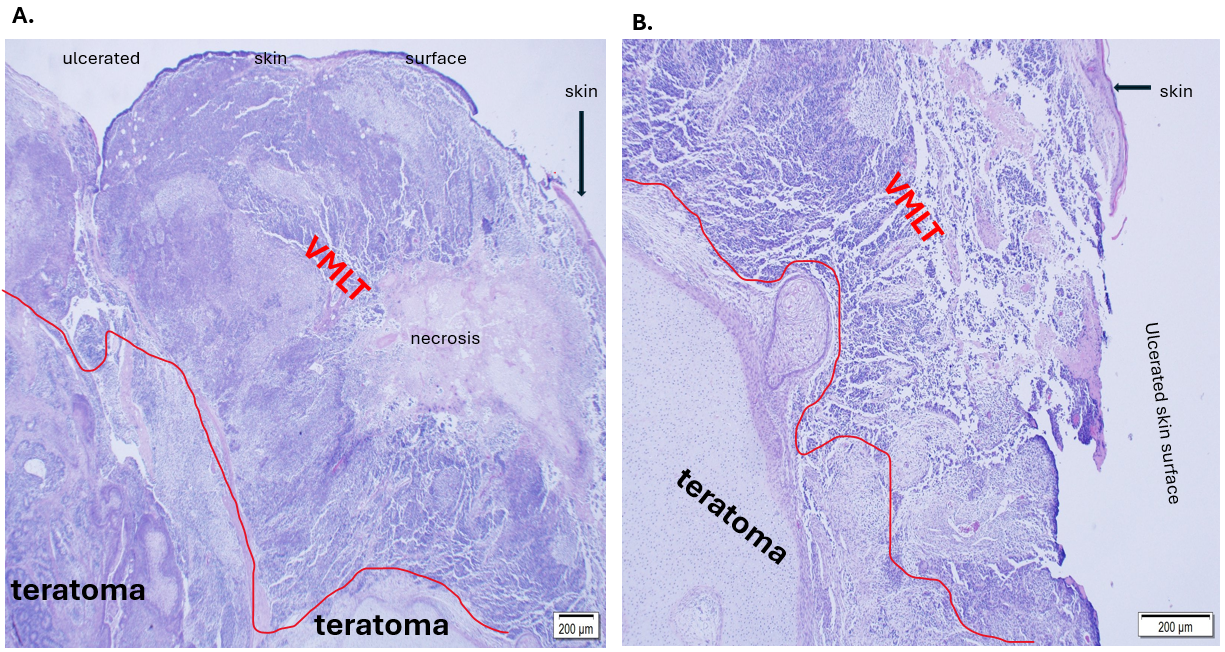

**Figure S7. Tumor T#85.3.** Large sheets of VMLT cells at skin surface covering deeper teratoma component in T#85.3 tumor (2X).

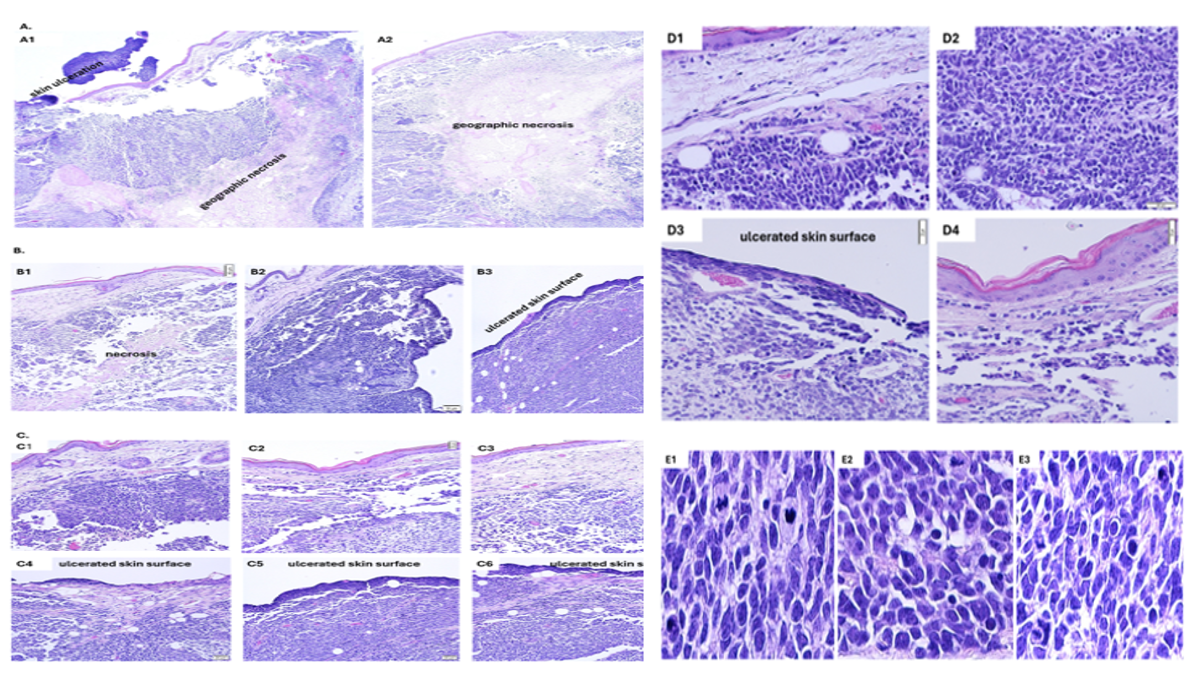

**Figure 8. Representative sections of VMLT in superficial mouse skin of T#85.3 with extensive necrosis and ulceration**. **(A)** 2X **(B)** 10X **(C)** 20X **(D)** 40X **(E)** HPF views of cytomorphology of VMLT cells like those of T#85.4 in Figure S5 (E).

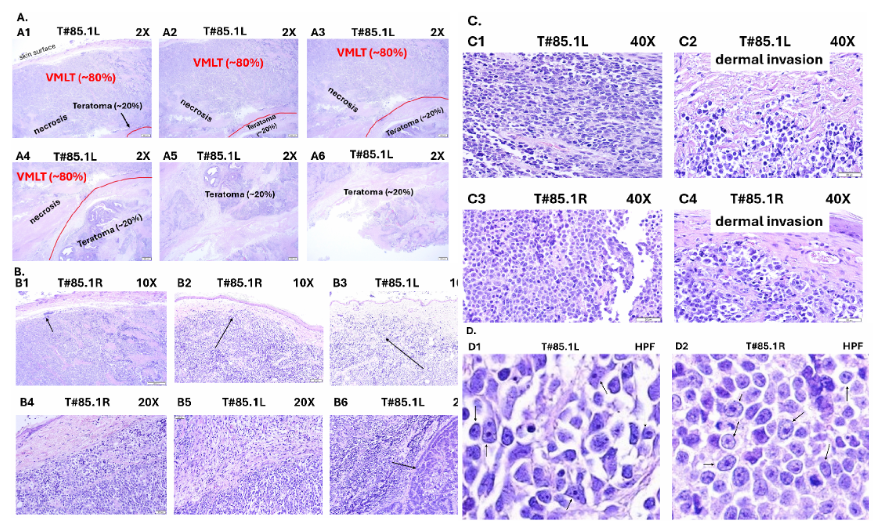

**Figure S9.** **Representative sections of VMLTs of T#85.1L & T#85.1R.**

A. Serial sections of T#85.1L to show extensive VMLT with necrosis in superficial mouse skin to deeper minor teratoma component at 2X
B. Sections of VMLT of tumor T#85.1L at 10X and 20X showed LVI (black arrow in B1), dermal invasion (B4, B5 and black arrows in B2 and B3) and sheets of small blue cells with rare foci of trabecular growth pattern (black arrow in B6).
C. Sections of VMLT of both T#85.1L and T#85.1R tumors at 40X showed sheets of small blue cells (C1 and C3 respectively) and dermal invasion of VMTL cells toward skin surface.
D. High power field views showed cytomorphology of VMLT cells of T#85.1L (D1) and T#85.1R (D2). Besides typical VP-MCC like cells with salt and pepper chromatin and inconspicuous nucleoli, intermixed larger atypical cells with paler vesicular chromatin, prominent large nucleoli and more abundant cytoplasm (black arrows) were also identified.

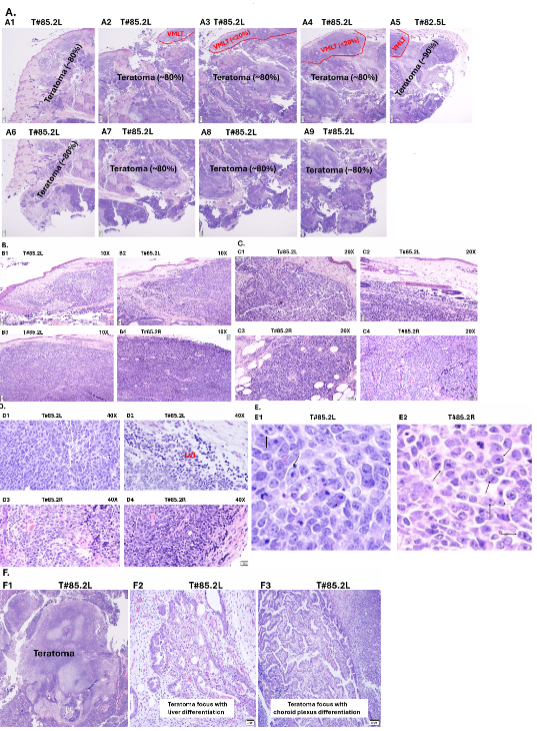

**Figure S10.** **Representative sections of VMLTs of T#85.2L and T#85.2R.** **(A)** Serial sections of entire T#85.2L tumor show a minor incipient VMLT component (~20% of tumor) at right superficial mouse skin and vast majority of teratoma component (~80%) at left and deeper (2X). Sections of VMLT of tumor T#85.2L and

T#85.2R at **(B)** 10X **(C)** 20X and **(D)** 40X show sheets of VMLT cells at superficial mouse skin with extensive

dermal invasion (B1-B4, C1-C3, D1 and D4) and LVI (D2). **(E)** High power field views show cytomorphology

of VMLT cells of T#85.2L (E1) and T#85.2R (E2). As well as typical VP-MCC-like cells with salt and pepper

chromatin and inconspicuous nucleoli, intermixed larger atypical cells with paler vesicular chromatin,

prominent large nucleoli and more abundant cytoplasm (black arrows) were also identified. (F) Extensive

teratoma component with three germinal layer differentiation in T#85.2L.

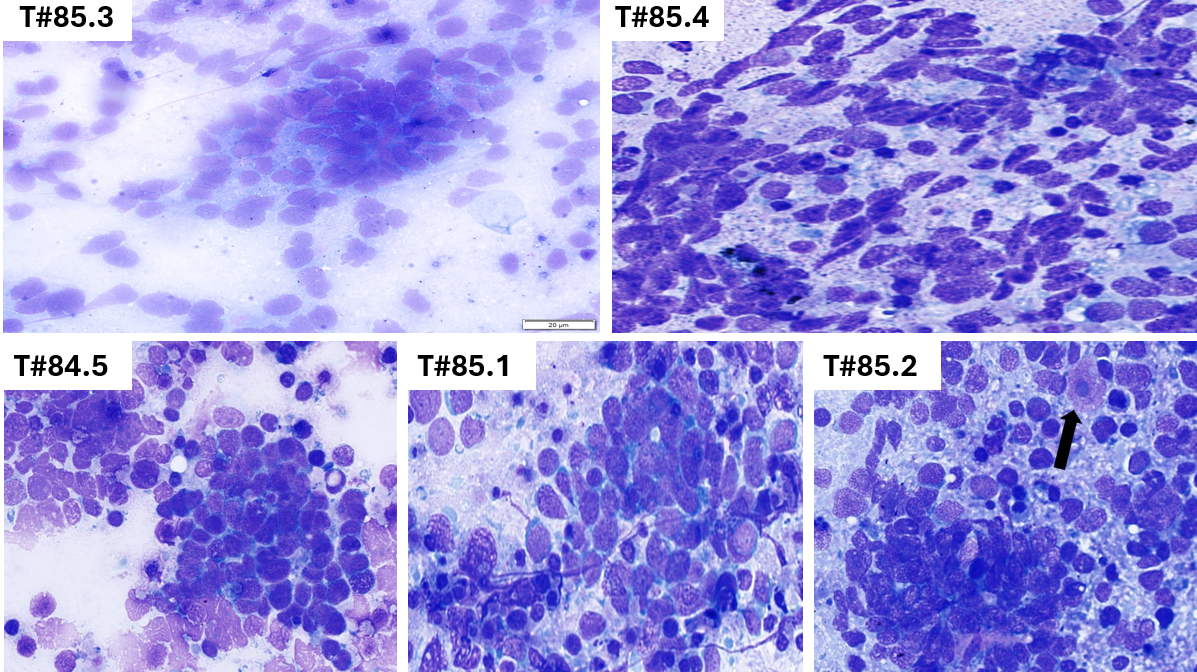

**Figure S11**. **Cytology smears of mouse CDX VMLT tumors.** T#85.4 and T#85.3 smears show typical SCNC cytomorphology with high N/C ratio, fine chromatin, inconspicuous nucleoli, nuclear streaking and nuclear molding. T#84.5, T#85.1 and T#85.2 show some cytomorphologic features atypical for VP-MCC: more abundant cytoplasm with some VMLT cells exhibiting prominent nucleoli (black arrow in T#85.2).

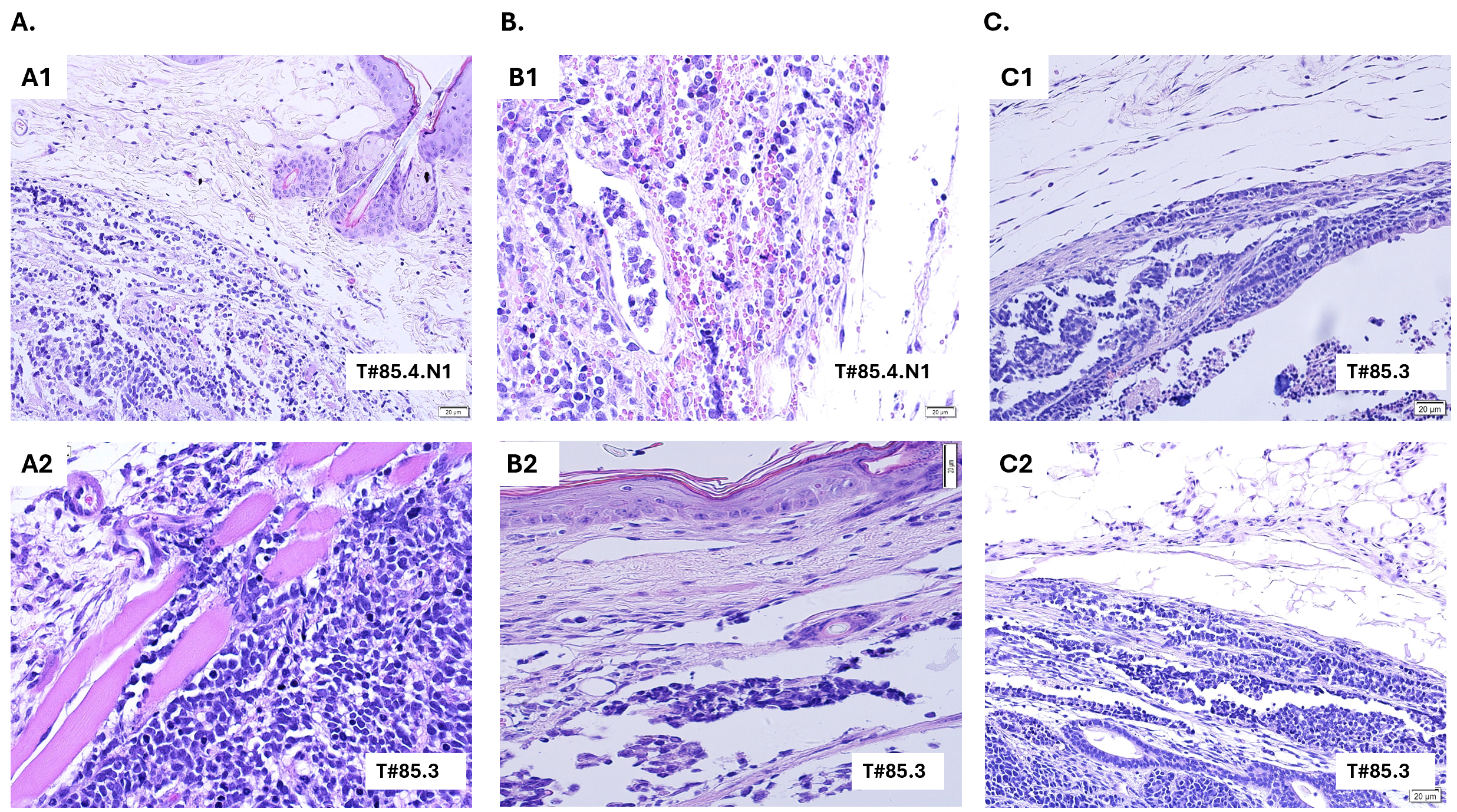

**Figure S12**. **VMLTs of T#85.3 and T#85.4 exhibit malignant histological features.** **(A)** Dermal invasion with VMLT cells infiltrating through dermal striated muscle layer in T#85.4 and T#85.3. **(B)** Lymphovascular invasion (LVI) with clusters of VMLT cells inside lymphatic spaces in T#85.4 and T#85.3. **(C)** Infiltration and destruction of tumor capsules by VMLT cells in T#85.4 and T#85.3.

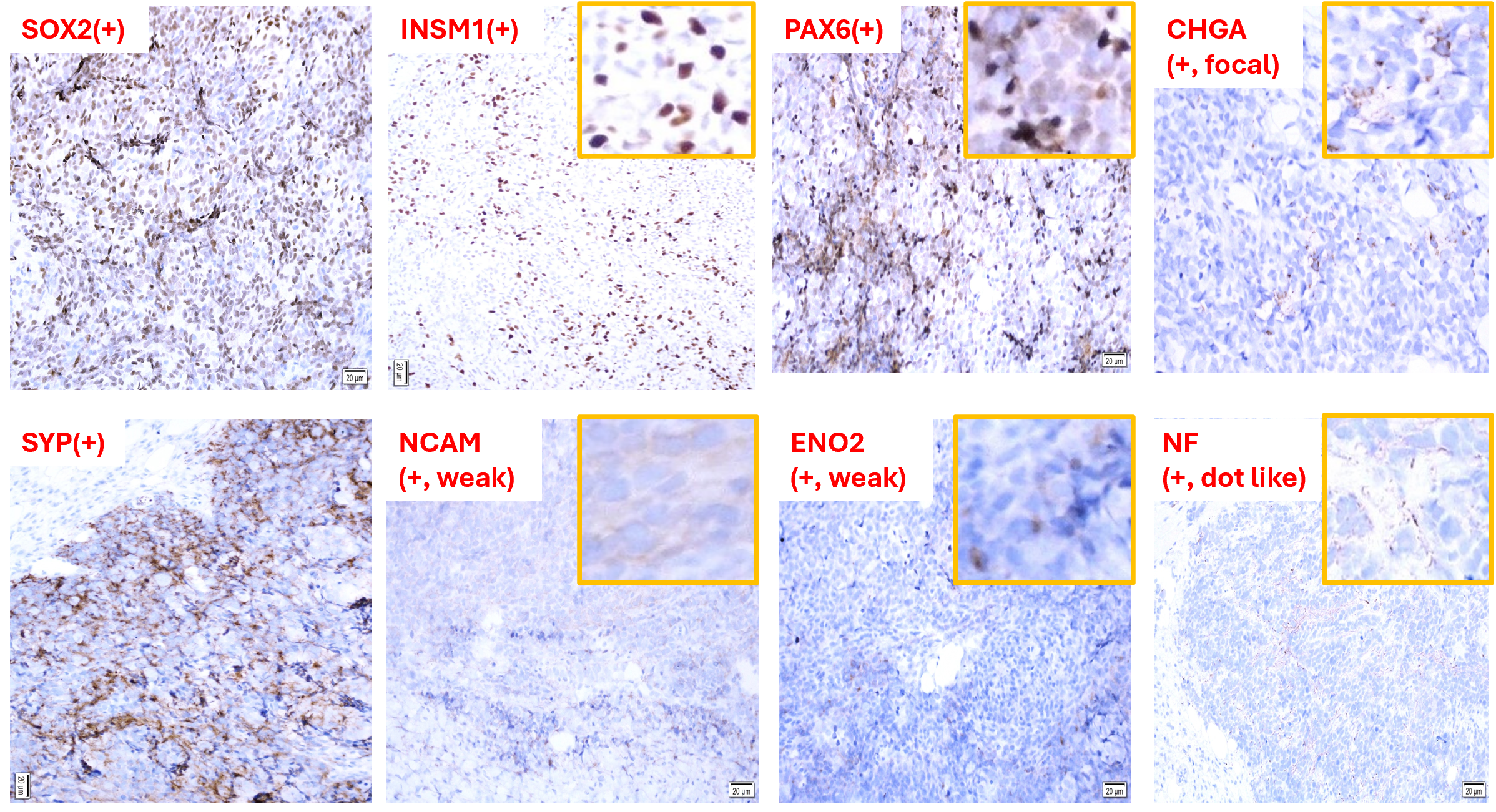

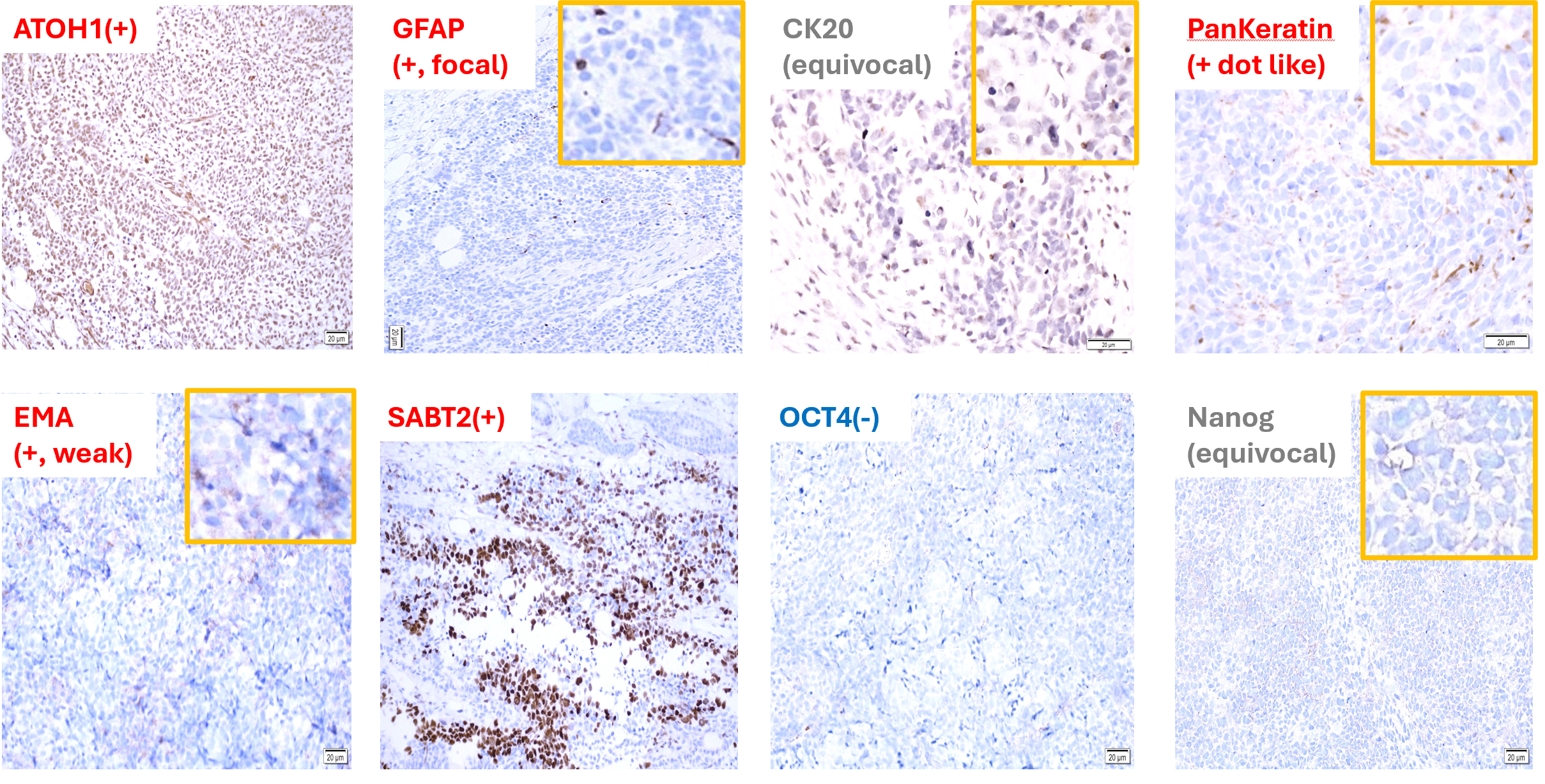

**Figure S13. Immunohistochemical stains of T#85.3.**

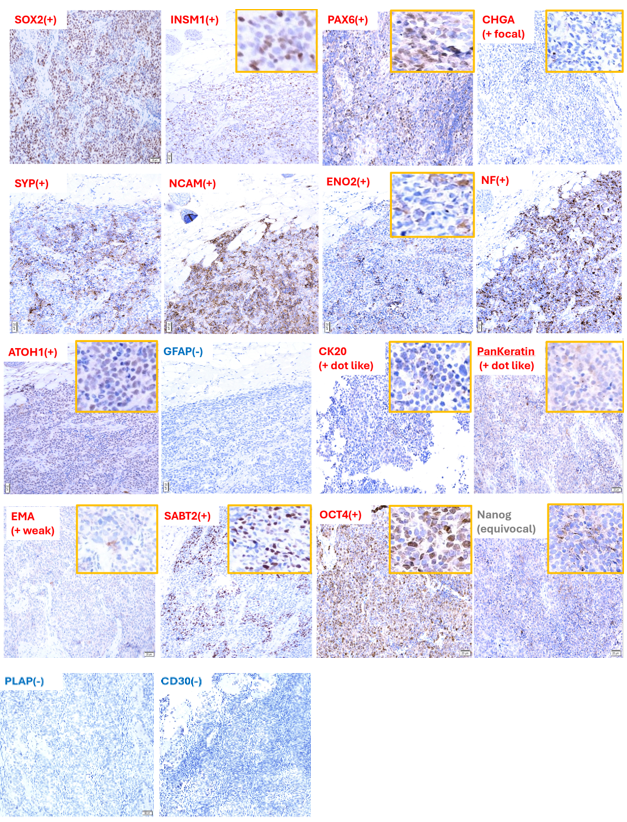

**Figure S14. Immunohistochemical stains of T#84.5R.**

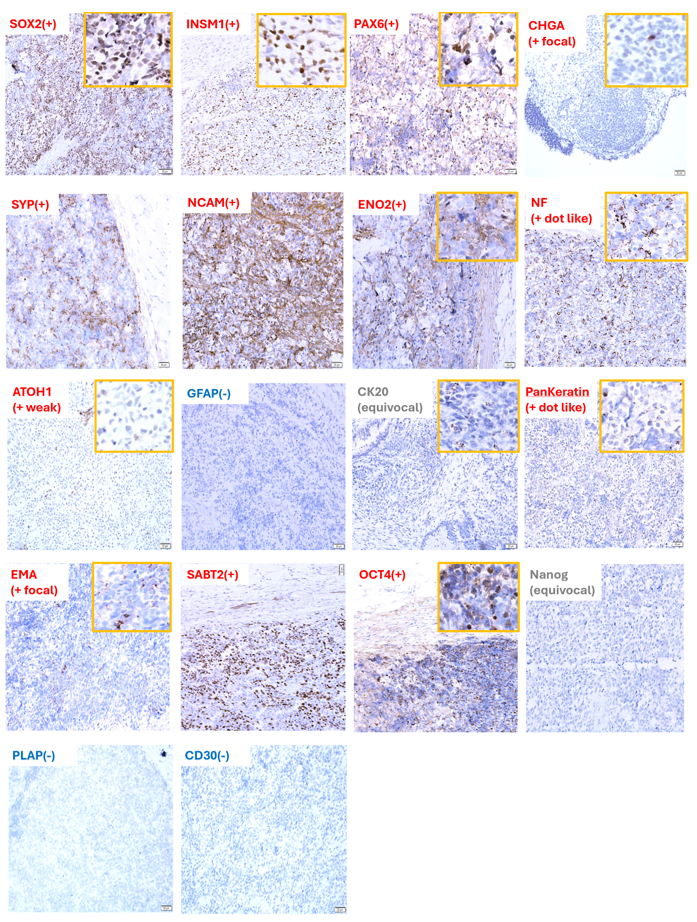

**Figure S15. Immunohistochemical stains of T#85.1L.**

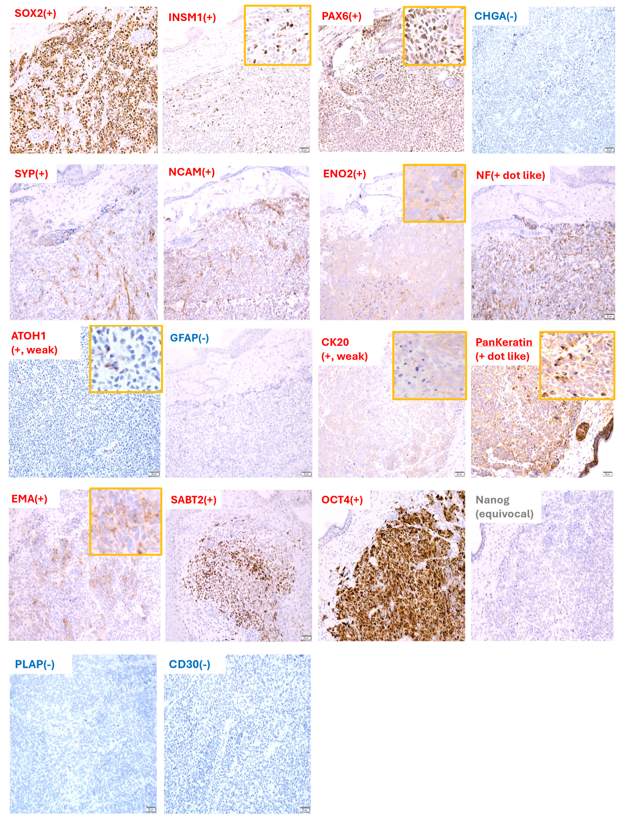

**Figure S16. Immunohistochemical stains of T#85.2L.**

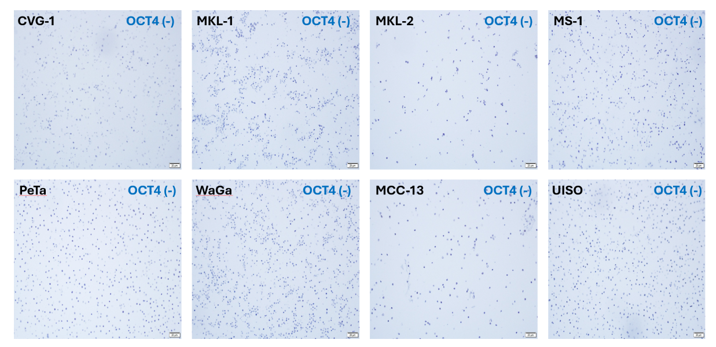

**Figure S17. Immunohistochemical stains of VP-MCC and VN-MCC cell lines.**

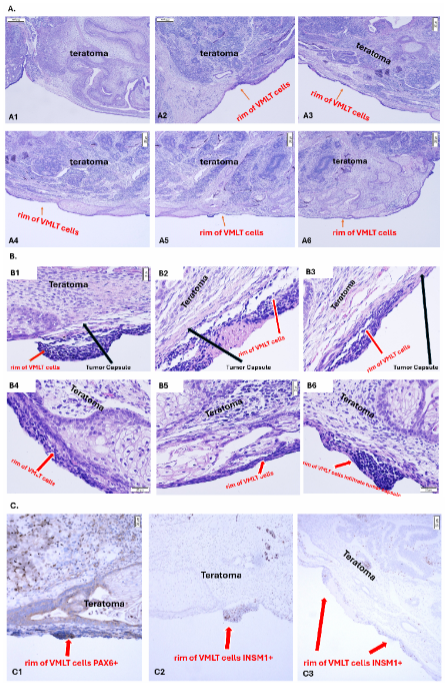

**Figure S18.** **Rim of VMLT cells infiltrate deep tumor capsule covering teratoma component with a differentiation gradient in T#85.3 tumor. (A)** Serial sections of the entire deep tumor capsular surface covering teratoma component above. A thin dark rim of VMLT cells can be seen lining the deep tumor capsule. A differentiation gradient can be seen in teratoma with the outer portion more mature. (2X). **(B)** HPF view of the dark rim shows it is indeed composed of VMLT cells. Differentiated epidermal epithelium and hair follicles are at the outermost part of the teratoma. **(C)** IHC staining shows the dark rim cells are INSM1(+) and PAX6(+), consistent with VMLT cell identity.

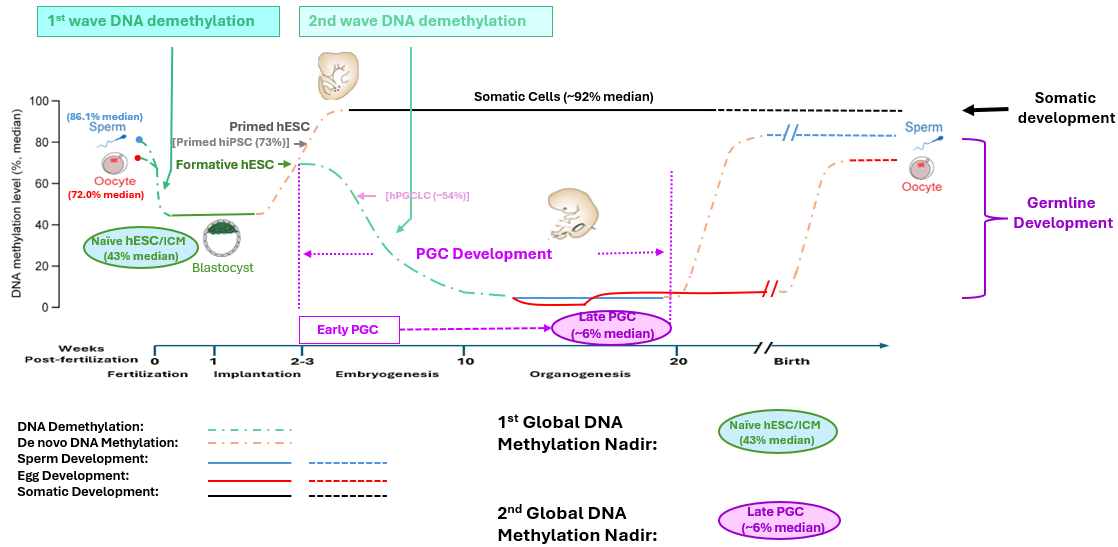

**Figure S19.** **Summary of prior study findings on two global DNA methylation (GDM) nadirs following two waves of global DNA demethylation during early embryogenesis and gametogenesis of human development** (Kobayashi et al., 2022; Wen et al., 2019). The 1^st^ wave of global DNA demethylation during the 1^st^ week of embryogenesis starts at the methylation level of the zygote after fertilization and decreases to the lowest level of naive hESCs/Inner Cell Mass (ICM) at the early blastocyst state, the 1^st^ GDM nadir (~43% median). Subsequently, de novo methylation starts and reaches the formative hESC state which will bifurcate to either the somatic fate or the germline fate. The somatic fate trajectory entails continued de novo methylation from the formative hESC state to the primed hESC state (mimicked by primed hiPSC ~73%) followed by somatic lineage differentiation until they develop into mature somatic cells with stable and high methylation levels (~92% median). The germline fate trajectory entails germline-specific 2^nd^ wave global DNA demethylation which starts when hPGCs are specified from formative hESCs with hPGCs becoming progressively demethylated along the hPGC developmental trajectory from early hPGCs (mimicked by hPGCLC ~54%) to late hPGCs, the 2^nd^ GDM nadir (~6% median). Subsequently, de novo methylation starts along the gonadal gametogenesis trajectory with GDM of developing germ cells gradually increasing until fertilization (sperm ~86.1% and oocyte ~72.0%). Adapted and used with permission from Wen et al., 2019.

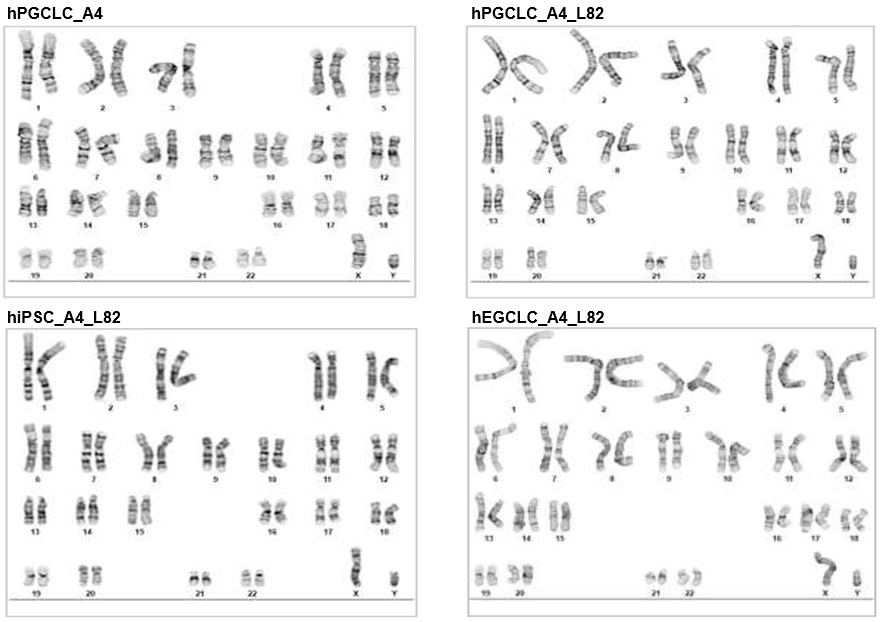

**Figure S20. Normal karyotype (46, XY) for all 4 primeval stem cell lines injected for the mouse study.**

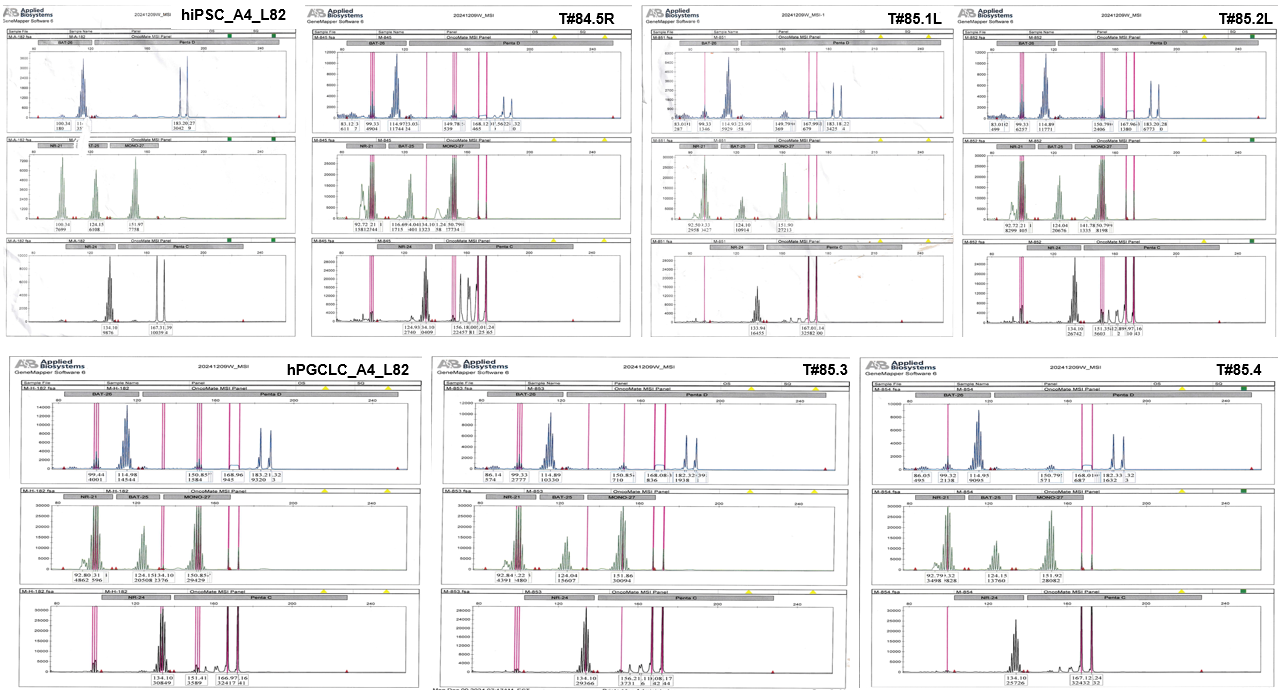

**Figure S21. Promega OncoMate MSI Analysis System was used for authentication of injected primeval stem cell lines and derived VMLT(+) tumors.** hiPSC_A4_L82 and derived VMLT(+) tumors T#84.5R, T#85.1L and T#85.2L as well as hPGCLC_A4_L82 and its derived VMLT(+) tumors T#85.3 and T#85.4 exhibit the same STR patterns including 5 mononucleotide repeats (NR-21, BAT-26, BAT-25, NR-24, MONO-27) and two dinucleotide repeats (Penta C, and Penta D).

**Tables S1, S3, S4, S5, S6:**

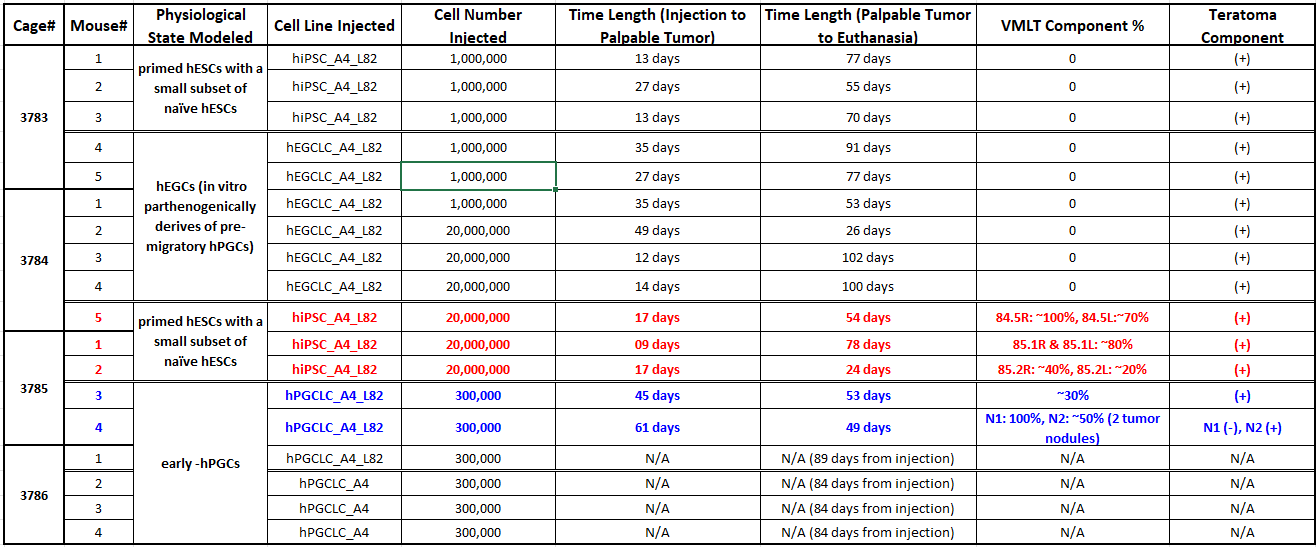

**Table S1. CDX Mouse Study Summary**

Table S2. (separate file)

**RT-qPCR Results of hPGCLC_A4_L82, hiPSC_A4_L82, Seven VMLTs, VP-MCC cell lines and VN-MCC cell lines**

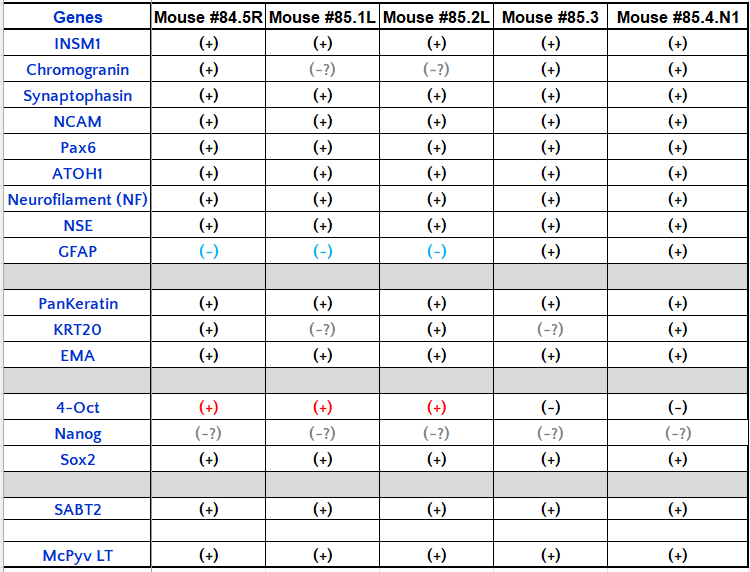

**Table S3. Results Summary of IHC Study of VMLTs**

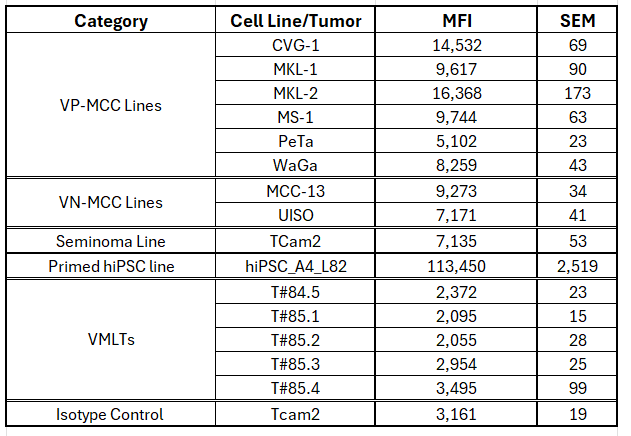

**Table S4. Global 5mC levels by flow cytometry. Median Fluorescence Intensity (MFI) and**

**Standard Error of the Mean (SEM) were calculated by the FlowJo software.**

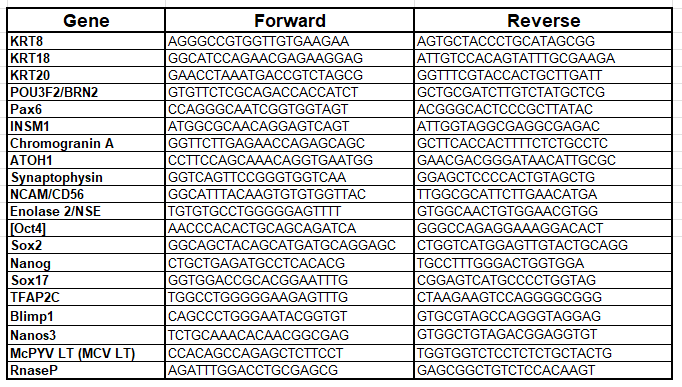

**Table S5. Primers for RT-qPCR**

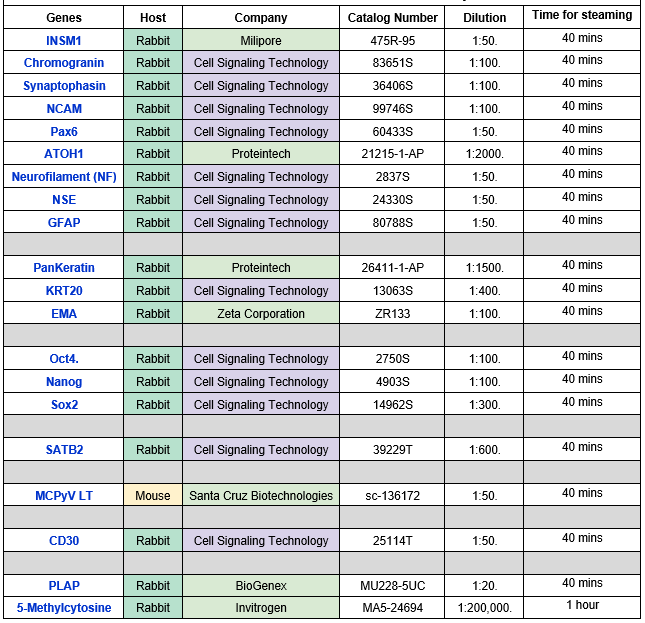
 **Table S6. Primary antibodies used for IHC studies.**
